# Structural basis for prohibitin-mediated regulation of mitochondrial *m*-AAA protease

**DOI:** 10.1101/2025.04.15.648916

**Authors:** Dingyi Luo, Lvqin Zheng, Ming-Ao Lu, Ziyao Chen, Panpan Wang, Qiang Guo, Ning Gao

## Abstract

Mitochondrial function critically dependents on protein quality control systems, with the *m*-AAA protease plays a key role at the inner mitochondrial membrane (IMM). The evolutionarily conserved prohibitins (PHBs) are essential modulators of this protease across species, yet the molecular mechanisms remain unclear.

Here, we present the Cryo-EM structure of the *Chaetomium thermophilum* PHB (*Ct*PHB) complex, revealing a cage-like assembly composed of 11 copies of PHB1/PHB2 heterodimers. Electron microscopic and biochemical analyses suggest that *m*-AAA proteases are enclosed within the PHB complex through interactions mediated by their SPFH-interacting motif (SIM) exposed in the intermembrane space. Further *in situ* cryo-ET directly visualizes these cage-protease assemblies in native mitochondria. Disruption of their interface leads to elevated *m*-AAA protease activity and diminished mitochondrial stress resistance. These data establish PHB complexes as spatial organizers that compartmentalize *m*-AAA proteases in membrane microdomains to fine tune proteolytic homeostasis.

Our findings reveal the critical role of the PHB complex in maintaining mitochondrial proteostasis, providing a unified mechanistic model to explain and reconcile the pleiotropic, and often contradictive phenotypes of PHBs and *m*-AAA protease in mitochondrial physiology and various disease conditions.

## Introduction

Mitochondria, the powerhouses of eukaryotic cells, orchestrate energy production, metabolic regulation, and cellular signaling^1^. Maintaining their functional integrity requires a multilayered protein quality control (PQC) system that coordinates protein folding, complex assembly, and targeted degradation of damaged components^2^. Dysregulation of mitochondrial PQC mechanisms leads to severe pathological consequences, particularly in neurodegenerative disorders^3,4^. The *m*-AAA (matrix-oriented ATPases associated with diverse cellular activities) protease, located in the inner mitochondrial membrane (IMM), plays a crucial role in mitochondrial PQC by degrading misfolded or damaged proteins^5^. Beyond its surveillance function, the *m*-AAA protease also processes precursor proteins, such as cytochrome c peroxidase (Ccp1)^6^ and mitochondrial ribosomal protein MRPL32^7^, ensuring their functional maturation. This dual function highlights its physiological importance and underscores the need for precise regulation, as both insufficient and excessive protease activity can severely disrupt mitochondrial homeostasis^8,9^.

The precise regulation of the *m*-AAA protease has long been associated to prohibitins (PHBs)^10^. PHBs, comprising two highly homologous proteins PHB1 and PHB2, belong to the SPFH (stomatin, prohibitin, flotillin, HflK/C) protein family^11^ and assemble into ring-shaped complex within the inner mitochondrial membrane (IMM)^12^, where they regulate the cristae morphogenesis^13^, and participate in the maintenance of respiratory chain complex integrity^14^ and mitochondrial DNA stability^15^. These functions highlight PHBs’ crucial role in maintaining both structural and functional integrity of mitochondria, which is essential for embryonic development as evidenced by the lethality of PHB-null mice^13^. Notably, the reduced PHB expression has been observed in specific brain regions of patients with neurodegenerative disorders, including the olfactory bulb in Alzheimer’s disease (AD)^16^ and the substantia nigra in Parkinson’s disease (PD)^17^. Meanwhile, PHBs are upregulated in various cancers^18–21^, where cells experience enhanced oxidative stress within the tumor microenvironment^22^. Presumably, the pleiotropic functions of PHB complex converge on their ability to regulate the broad substrate spectrum of *m*-AAA protease ^23–25^. The intimate functional relationship between the PHB complex and *m*-AAA protease is evidenced by their conserved physical association across species, from yeast^7,26^ to mammalian cells^27,28^, and further supported by their coordinated expression patterns during metabolic shifts and oxidative stress ^29,30^. Collectively, PHBs emerge as central regulators safeguarding IMM proteostasis and membrane dynamics through their integrated control of *m*-AAA proteases, which execute dual functions in mitochondrial protein quality control and processing.

Structural studies of prokaryotic PHB*-m*-AAA protease homologs reveal that HflK/C complexes form a caged structure around FtsH protease, creating a barrier between its catalytic center and membrane protein substrates^31,32^. However, whether this regulatory mechanism is conserved in eukaryotic mitochondria remains unclear. Unlike their bacterial counterparts, eukaryotic PHBs also undergo dynamic phosphorylation, which has been implicated in modulating cell death ^33^ and stress responses during immune signaling^34^. This post-translational modification likely represents an additional layer of regulation for mitochondrial function. Consequently, the molecular mechanisms underlying PHB-mediated control of *m*-AAA protease activity and its role in maintaining mitochondrial homeostasis in eukaryotes remain to be fully elucidated.

Here, we explored the molecular basis of PHB-mediated mitochondrial proteostasis. Cryo-EM analysis revealed that the *Chaetomium thermophilum* PHB complex forms a cage-like assembly, establishing specialized membrane microdomains within the IMM. Phosphomimetic mutations disrupted the PHB assembly, suggesting that post-translational modification is another layer of this regulatory system. Structural and functional analysis demonstrated that PHB cages enclose the *m*-AAA protease to spatially confine its proteolytic activity, and cryo-ET visualization confirmed this regulatory architecture in native IMM. Together, our findings propose a model for how cells control mitochondrial proteostasis through the PHB-*m*-AAA protease supercomplex.

## Results

### *Ct*PHB complex forms a membrane attached 22-mer assembly

To elucidate the molecular mechanisms underlying PHB complex function, we performed structural characterization of PHB complexes from several model organisms. Using *Saccharomyces cerevisiae* as the expression host, we produced proteins of PHB homologs from multiple species, including *Homo sapiens* (*Hs*PHBs), *Saccharomyces cerevisiae* (*Sc*PHBs) and *Chaetomium thermophilum* (*Ct*PHBs) (Fig. S1–2). The *Ct*PHB complex were selected for structural studies due to their high sample homogeneity. Initial three-dimensional classification of the cryo-EM data revealed an unanticipated density corresponding to the yeast mitochondrial chaperone Hsp60 (Fig. S3a, b). This association was deemed non-physiological given that Hsp60, a soluble matrix protein, would be spatially segregated from the membrane-embedded PHB complex under native conditions. Consistent with this interpretation, the interaction was abolished under mild detergent conditions during purification (Fig. S3c, d), confirming its nature as a purification artifact. Following computational subtraction of the Hsp60 density and subsequent 3D classification, initial C1 refinement at 8.7 Å resolution revealed an apparent 22-fold symmetry, indicating that a total of 22 PHB subunits form a circular complex. Bacterial HflK/C alternate to form a 24-mer complex ^31,32^, and an alternating arrangement of PHB1 and PHB2 subunits within the PHB complex was also hypothesized ^35^. Therefore, we performed C11 symmetry-constrained refinement that yielded the high-resolution structure of the *Ct*PHB complex at 3.2 Å (Fig. S4, 5), which confirmed this alternative subunit arrangement, and the map quality is sufficient to unambiguously assign PHB1/PHB2 subunits.

Structural analysis elucidates a precisely organized cage-shaped architecture anchored within the inner mitochondrial membrane (IMM), with the cage projecting approximately 90 Å into the intermembrane space (IMS) (Fig. 1a, b). The molecular assembly exhibits three distinct architectural parts: a membrane-proximal region comprising transmembrane helices and SPFH1 domain segments integrated into the IMM; a central structural element extending 55 Å in height, composed predominantly of SPFH domains that establish a rigid scaffold through extensive inter-subunit interactions; and an upper region spanning 35 Å in height, consisting of Coiled-coil (CC) domains that generate the characteristic cap-like topology. When viewed along the membrane normal, the complex reveals a sophisticated oligomeric arrangement comprising 11 copies of *Ct*Phb1 and 11 copies of *Ct*Phb2 subunits in precise alternation, forming a highly ordered circular assembly, with the diameters at the membrane-proximal and distal regions being 180 Å and 100 Å, respectively. Notably, the interior of the cage is freely accessible through the central channel in the cap (approximately 25 Å in diameter) (Fig. 1c–f).

**Figure 1.**
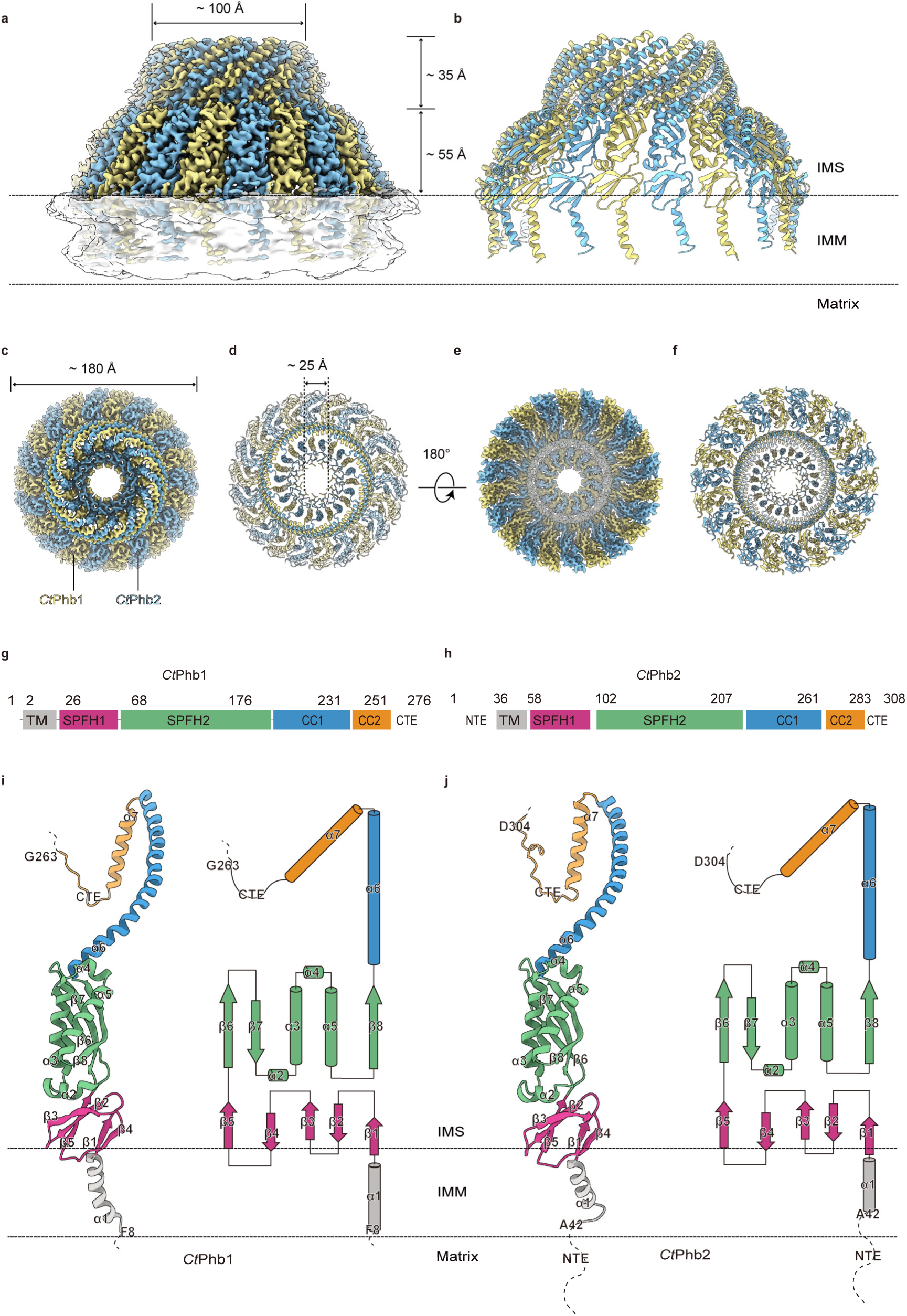
Cryo-EM Structure and Domain Organization of the *Ct*PHB Complex. **a, b,** Side views of the cryo-EM density map (a) and cartoon representation (b) of the *Ct*PHB complex. The cage-like structure projects ∼90 Å into the IMS, with a membrane-proximal region, a central body (∼55 Å) and an upper coiled-coil region (∼35 Å). *Ct*Phb1 and *Ct*Phb2 are shown in yellow and blue, respectively. **c–f,** Top (c, d) and bottom (e, f) views showing the 11-fold symmetric assembly with ∼180 Å diameter and a central pore of ∼25 Å. **g, h,** Domain architecture of *Ct*Phb1 (g) and *Ct*Phb2 (h). Domains are colored as follows: transmembrane helix (gray), SPFH1 (magenta), SPFH2 (green), CC1 (blue) and CC2(orange). Numbers indicate domain boundaries. **i, j,** Structural organization and topology diagrams of *Ct*Phb1 (i) and *Ct*Phb2 (j) protomers showing secondary structure elements relative to mitochondrial compartments. Dashed lines represent unresolved regions, including the matrix-facing NTE of *Ct*Phb2.

Comprehensive structural analysis demonstrates that both *Ct*Phb1 (276 residues) and *Ct*Phb2 (308 residues) exhibit conserved domain architectures, comprising an N-terminal transmembrane helix (TM), followed by tandem SPFH domains (SPFH1 and SPFH2), dual CC regions (CC1 and CC2), and terminating in a flexible C-terminal extension (CTE) (Fig. 1g, h). The atomic model encompasses residues 8–263 of *Ct*Phb1 and residues 42–304 of *Ct*Phb2, with the N-terminal residues 1–41 of *Ct*Phb2 presumably constituting a flexible extension within the matrix space. The SPFH1 and SPFH2 domains of both *Ct*Phb1 and *Ct*Phb2 maintain highly conserved structural folds, similar to other SPFH family proteins^31,32,36,37^ (Fig. 1i, j). The CC regions adopt extended α-helical conformations that facilitate critical inter-subunit interactions essential for complex assembly. The C-terminal tails exhibit conformational heterogeneity, precluding their visualization in the final structure.

### Structural basis for PHB-mediated membrane microdomain organization

The *Ct*PHB complex is stabilized by extensive inter-subunit interactions spanning from the SPFH domains to the CC regions (Fig. S6). Consistent with the essential role of CC regions in complex assembly, as demonstrated by previous deletion studies^12^, the parallel arrangement of CC1 and CC2 regions between adjacent subunits forms a distinctive cap-like structure at the complex apex (Fig. 2a). At the CC1-CC2 junction, an intricate network of inter-subunit or intra-subunit hydrophobic and polar interactions stabilizes this architectural element (Fig. 2b). A highly conserved tyrosine residue (Y266) in *Ct*Phb2 appear to function as a central organizing element within this interface. Its hydroxyl group lies 2.9 Å from the carboxylate group of E217 in the adjacent *Ct*Phb1, while its aromatic ring participated in hydrophobic interactions with surrounding residues (Fig. 2c), establishing the characteristic fold that determines the precise positioning of these helical segments. This tyrosine residue in *Ct*Phb2 is evolutionarily conserved (Fig. S7), with the corresponding residue Y248 in *Hs*PHB2 serving as a critical regulatory phosphorylation site^34^, highlighting the fundamental importance of this position in PHB complex regulation.

**Figure 2.**
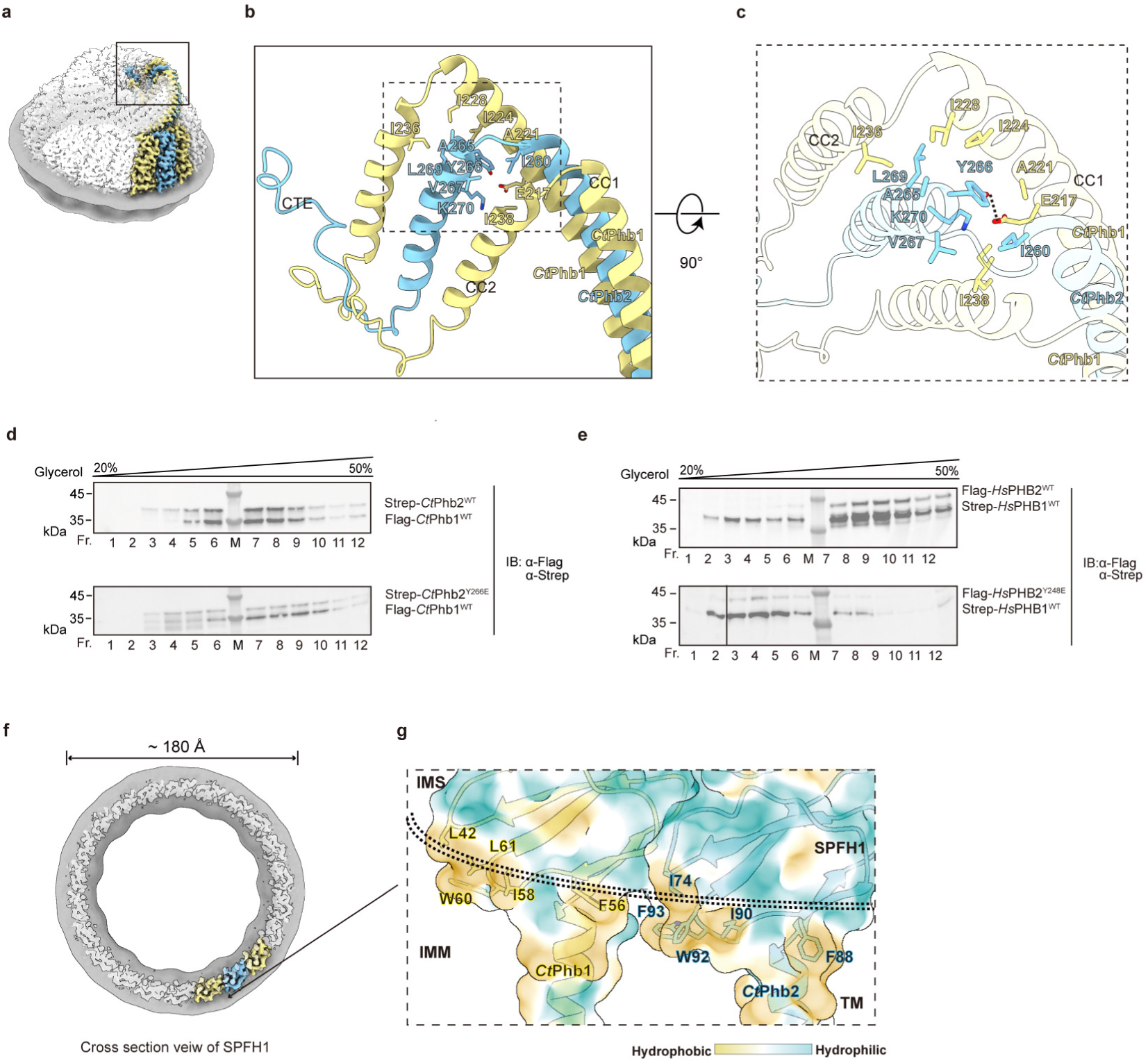
Structural Organization of the *Ct*PHB Complex Reveals Critical Determinants for Assembly and Membrane Domain Formation. **a–c,** Structure of the coiled-coil region interface. Overview of the complex highlighting the CC region (a), detailed view of CC1-CC2 junction (b), and close-up view after 90° rotation showing the detailed interaction network (c; the hydrogen bond shown as a dashed line). **d, e,** Glycerol gradient analysis (20**–**50%) showing complex destabilization by phosphomimetic mutations. Wild-type complexes concentrate in high molecular weight fractions, while *Ct*Phb2^Y266E^ (d) and *Hs*PHB2^Y248E^ (e) mutations cause complex disassembly, as detected by immunoblotting with anti-Flag and anti-Strep antibodies (Fr., fraction; M, marker). **f, g,** Membrane microdomain organization. Cross-sectional view showing the ∼180 Å diameter membrane domain (f) and surface hydrophobicity analysis revealing conserved membrane-interacting residues in the SPFH1 domains (g). Hydrophobicity scale: yellow (hydrophobic) to blue (hydrophilic).

To experimentally assess the structural and functional importance of Y266 in *Ct*Phb2, we introduced a negative charge at this position through Y266E mutation to examine its effects on complex assembly. Glycerol gradient analysis revealed that wild-type *Ct*PHB complex exhibited sharp, concentrated distribution in high molecular weight fractions (Fr. 6-8), whereas the *Ct*Phb2^Y266E^ mutant resulted in loss of the distinct *Ct*Phb2 peak and appearance of degradation products in lower molecular weight fractions (Fig. 2d). This destabilization suggested that introducing a negative charge at the critical corner region of the CC domain compromises complex integrity. Notably, mutation of the corresponding residue in human PHB2 (*Hs*PHB2^Y248E^) showed even more pronounced effects on complex stability (Fig. 2e). Guided by structural predictions and analysis of reported phosphorylation sites, we further examined additional corner residues within the CC domain of *Hs*PHB1. Phosphomimetic mutations at these positions (*Hs*PHB1^Y259E^, *Hs*PHB1^S265E^) also demonstrated varying degrees of complex destabilization (Fig. S8b), suggesting that post-translational modifications at these conserved sites could serve as regulatory mechanisms for complex integrity.

When viewed in cross-section parallel to the membrane plane, the precisely organized membrane-associated regions of the properly assembled complex create a defined membrane microdomain approximately 180 Å in diameter (Fig. 2f). This microdomain is enclosed by the SPFH1 domains, which contain multiple conserved hydrophobic residues that insert into the IMM (Fig. 2g). While these membrane-interacting residues show sequence variation across species (Fig. S7), they maintain their hydrophobic nature, reflecting evolutionary adaptation to diverse membrane environments. This circular organization and membrane-inserting ability of the SPFH1 domains are also seen in other SPFH proteins^31,32,36,37^, emphasizing that the membrane microdomains established by the PHB complex within the IMM^38^ is the structural foundation for its function in mitochondrial physiology

### Conserved structural basis of PHB-*m*-AAA protease interaction

To directly visualize the interaction between PHB complexes and *m*-AAA proteases, we co-expressed the *Ct*PHB complex components (*Ct*Phb1 and *Ct*Phb2) together with its *m*-AAA protease *Ct*Rca1 (Respiratory chain assembly 1) through yeast expression system. Initial nsEM analysis of purified *Ct*Rca1 alone revealed well-assembled particles with characteristic double-layered architecture, comprising stacked ATPase and protease domains, with adjacent density corresponding to the detergent-solubilized transmembrane regions and intermembrane space domain (IMSD) (Fig. S9). Building on our observation that DDM extraction led to artificial Hsp60 association with *Ct*PHB complex, we systematically optimized the sample preparation conditions using milder detergents (digitonin), along with refined growth media and cell disruption methods (see methods for details). Under these optimized conditions, we successfully co-purified the *Ct*PHB-*Ct*Rca1 complex. Subsequent nsEM analysis and 2D classification revealed an architectural arrangement where the PHB complex forms a cage-like structure that encapsulates the *m*-AAA protease (Figure 3a, b). Similarly, nsEM analysis of co-purified *Homo sapiens* PHB-*m*-AAA protease complexes showed characteristic particles with comparable cage-protease organization (Fig. S8c, d), suggesting a conserved structural arrangement across species. These structural analyses clearly revealed that the PHB complex forms a cage-like structure around the *m*-AAA protease, with interaction occurring through the IMSD region (Fig. 3c).

**Figure 3.**
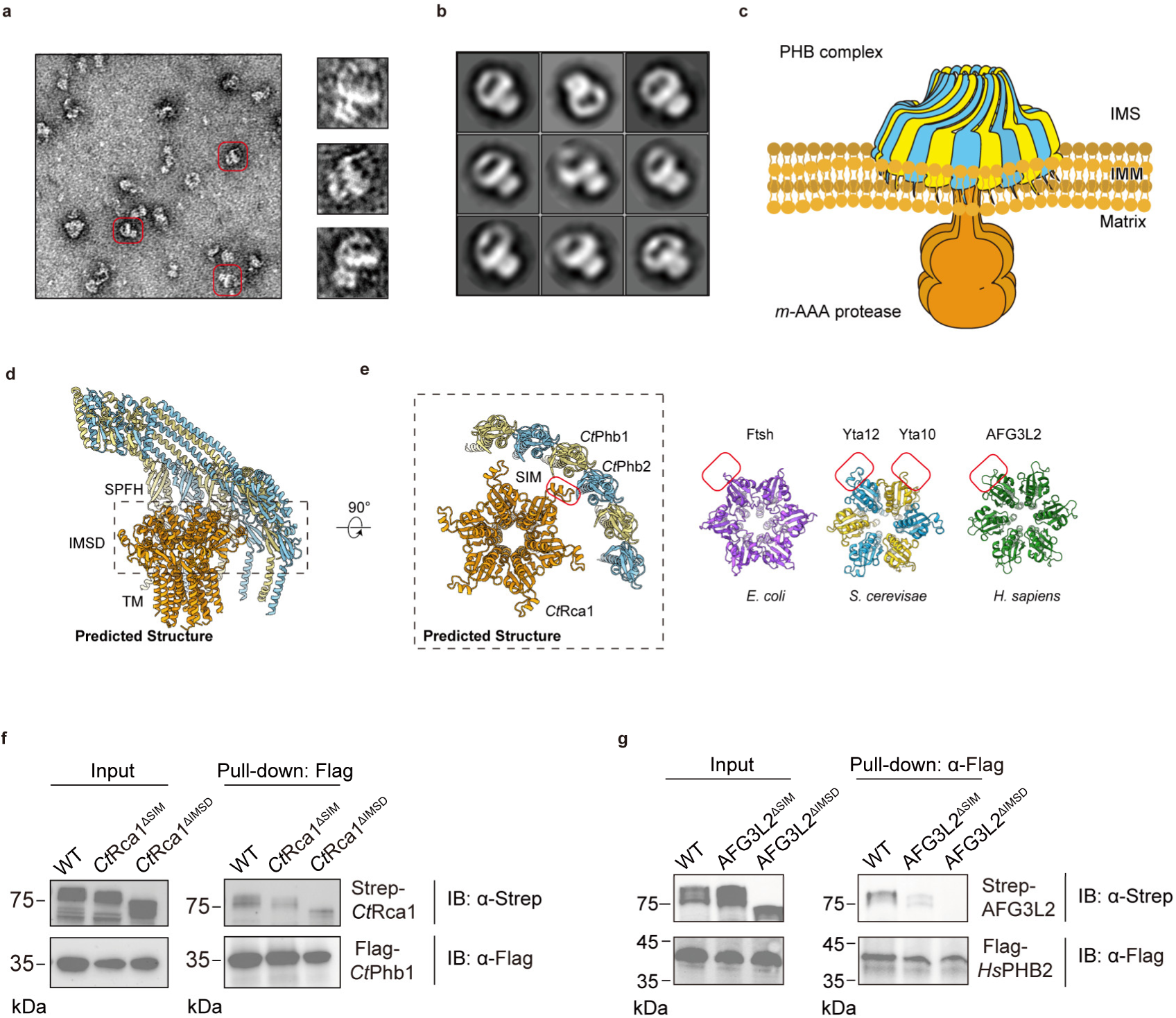
Structural and biochemical characterization of PHB-*m*-AAA protease interaction. **a, b,** nsEM analysis of the *Ct*PHB-*Ct*Rca1 supercomplex. Representative micrograph with selected particles (red boxes) and individual particle images (a). Reference-free 2D class averages showing characteristic cage-protease organization (b). **c,** Schematic illustration of PHB complex (blue/yellow) encapsulating *m*-AAA protease (orange) in the inner mitochondrial membrane (IMM). **d,** AlphaFold-predicted structure of the *Ct*PHB-*Ct*Rca1 complex. Side view showing SPFH and IMSD domains (left) and top view highlighting the interaction interface (dashed box, right). **e,** Top views of *m*-AAA proteases’ IMSD from different species, revealing spatially conserved SIM regions (red boxes). **f, g,** Pull-down assays validating SIM-mediated interactions between Strep-tagged *Ct*Rca1 and Flag-tagged *Ct*Phb1 (f), or Strep-tagged AFG3L2 and Flag-tagged *Hs*PHB2 (g), comparing wild-type and mutant variants by immunoblotting.

To investigate whether the observed cage-like architecture reflects a specific recognition mechanism rather than non-specific encapsulation, we performed AlphaFold prediction analysis, which revealed a potential interaction interface where *Ct*Rca1 contains an exposed motif that could mediate its association with the PHB complex (Figure 3d). Intriguingly, structural comparison across different species showed that while *m*-AAA proteases from *Escherichia coli* (FtsH) to *Homo sapiens* (AFG3L2) maintain similar overall architectures, these interaction motifs, which notably include the previously reported residues mediating FtsH-HflK/C interaction in *Escherichia coli*^31^, occupy structurally equivalent exposed positions despite showing limited sequence conservation (Fig. 3e, Fig. S8e, f). Given their specific localization and potential role in complex assembly, we designated these regions as SPFH-interacting motifs (SIMs). To validate the functional significance of SIMs, we strategically generated two variants: one lacking the entire IMSD and another specifically lacking the SIM. Pull-down experiments with both *Ct*PHB-*Ct*Rca1 and *Hs*PHB-AFG3L2 pairs revealed that while deletion of the IMSD abolished the interaction as expected, specific removal of the SIM alone was sufficient to disrupt association compared to the wild-type (Figure 3f, g). Together, these structural and functional analyses demonstrate a conserved interaction mechanism mediated by spatially preserved motifs with divergent sequences across species.

### PHB fine-tunes *m*-AAA protease activity in its functions of mitochondrial protein processing and stress tolerance

To investigate the physiological significance of the conserved interactions between the PHB complex and *m*-AAA protease, we examined how SIM-mediated association influences *m*-AAA protease activity and mitochondrial function. We engineered *Saccharomyces cerevisiae* strains lacking either the PHB complex components (*Δscphb1* and *Δscphb2*) or specific domains of the *m*-AAA protease subunit Yta12 (*yta12^ΔSIM^*and *yta12^ΔIMSD^*). Using MrpL32, a well-characterized *m*-AAA protease substrate, as a reporter in isolated mitochondria, we found that both *Δscphb1* and *yta12^ΔSIM^* mutations enhanced substrate processing, while *yta12^ΔIMSD^* led to substrate accumulation (Fig. 4a, b). The substrate accumulation in *yta12^ΔIMSD^* likely reflects compromised protease assembly, analogous to the established role of the corresponding membrane domain (comprising two transmembrane segments and the intervening periplasmic region) in bacterial FtsH oligomerization^39^. These results indicate that the PHB complex exerts an inhibitory effect on *m*-AAA protease activity towards its native substrate through SIM-mediated interactions.

**Figure 4.**
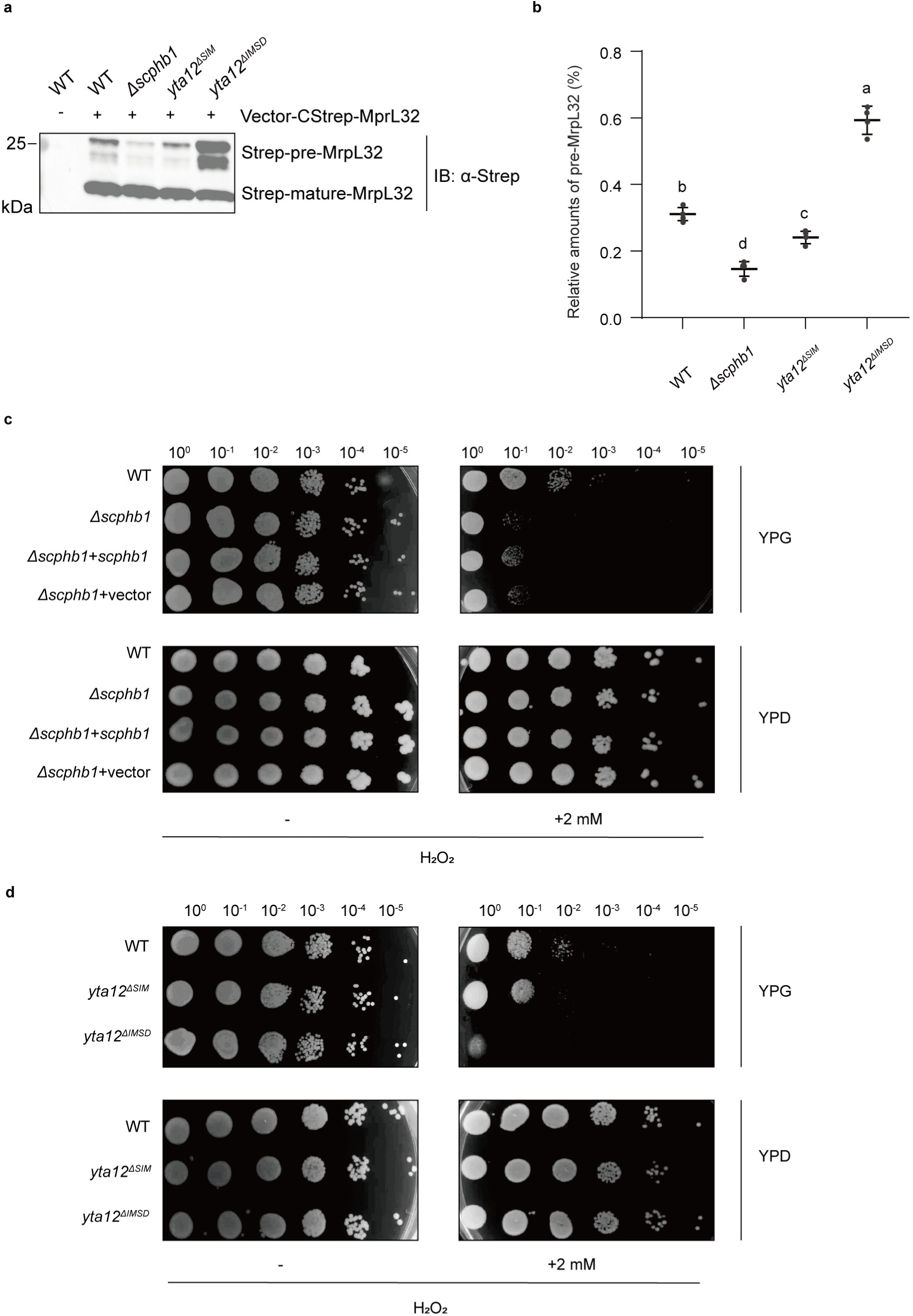
PHB complex regulation of *m*-AAA protease activity impacts oxidative stress tolerance. **a, b,** Analysis of MrpL32 processing in mitochondria. Immunoblot showing precursor and mature forms in wild-type and mutant strains (a). Quantification of precursor levels relative to total MrpL32 from four independent experiments; data are presented as mean ± S.D.; different letters indicate statistically significant differences between groups (*p* < 0.05, one-way ANOVA followed by Tukey’s test) (b). **c, d,** Oxidative stress sensitivity assays. Serial dilutions (10^0^-10^-5^) of indicated strains were spotted on YPG (respiratory) or YPD (fermentative) media ± 2 mM H_2_O_2_. Wild-type and *Δscphb1* strains with indicated complementation (c). Wild-type and yta12 domain deletion mutants (d). Images were taken after 3 d (YPG) or 2 d (YPD) at 30°C.

PHB deficiency in Kit225 cells was previously shown to enhance cellular sensitivity to H_2_O_2_-induced cell death, suggesting a protective role of the PHB complex against oxidative stress^18^. Extending these findings, we explored whether this function of the PHB complex is mediated through fine-tuning the activity of *m*-AAA protease. Using the yeast-based spot assay, we examined the fitness of PHB1/2 or *m*-AAA protease mutant cells in non-fermentable medium (YPG), where glycerol utilization requires intact mitochondrial metabolic function^40^. Firstly, a complete deletion of *yta12* abolished growth in YPG (Fig. S10a), consistent with established role of *m*-AAA protease in mitochondrial respiratory function ^41,42^. In contrast, *yta12^ΔSIM^* and *yta12^ΔIMSD^* strains were still viable in YPG, indicating that these mutants retained the capacity to maintain basal IMM protein homeostasis. Secondly, *Δscphb1* and *Δscphb2* strains both exhibited elevated sensitivity to H_2_O_2_-induced oxidative stress specifically in YPG medium, and this phenotype could be partially rescued by vector-based expression of *Sc*Phb1 or *Sc*Phb2 (Fig. 4c, Fig. S10b). Notably, the oxidative stress sensitivity was absent in fermentable YPD medium (Fig. 4c, Fig. S10b), where glucose metabolism primarily relies on cytosolic glycolysis, suggesting that loss of PHB specifically compromises mitochondrial stress resilience. Thirdly, *yta12^ΔSIM^* cells with presumably disrupted PHB-*m*-AAA protease interface displayed a similar hyper-sensitivity to oxidative stress in YPG medium as the *Δscphb1* and *Δscphb2* cells did (Fig. 4d), further demonstrating that SIM-mediated interaction is essential for mitochondrial stress response. Importantly, the levels of sensitivities of these yeast mutants correlated very well with the abilities of *Homo sapiens and Chaetomium thermophilum* homologs in forming stable PHB-*m*-AAA protease supercomplexes (Fig. 3f, g). On the other hand, *yta12^ΔIMSD^* cells with reduced proteolytic activity also rendered mitochondria more vulnerable to oxidative damage (Fig. 4d), indicating a minimal requirement of protease activity for proper stress tolerance. These findings suggest that an appropriate range of *m*-AAA protease activity, particularly through SIM-mediated PHB control, is essential for maintaining IMM protein homeostasis against oxidative stress (as detailed in Discussion).

### *In situ* visualization of conserved PHB-protease architecture in the IMM

To validate the molecular architecture of PHB and its ability to enclose *m*-AAA proteases in cells, we performed cryo-electron tomography on mitochondrial regions of intact HEK293T cells. Tomographic reconstructions revealed abundant cage-like molecular assemblies oriented toward the IMS, decorating the mitochondrial cristae membrane (Fig. 5a, b), reminiscent of our *in vitro* resolved *Ct*PHB structure. To identify these structures, we conducted subtomogram averaging analysis using 450 particles from 17 tomograms. The resulting density map showed cage-shaped assemblies matching the dimensions of our *Ct*PHB complex (Fig. 5c, d). Molecular fitting of atomic models confirmed structural conservation between isolated and membrane-embedded PHB complexes (Fig. 5c, d).

**Figure 5.**
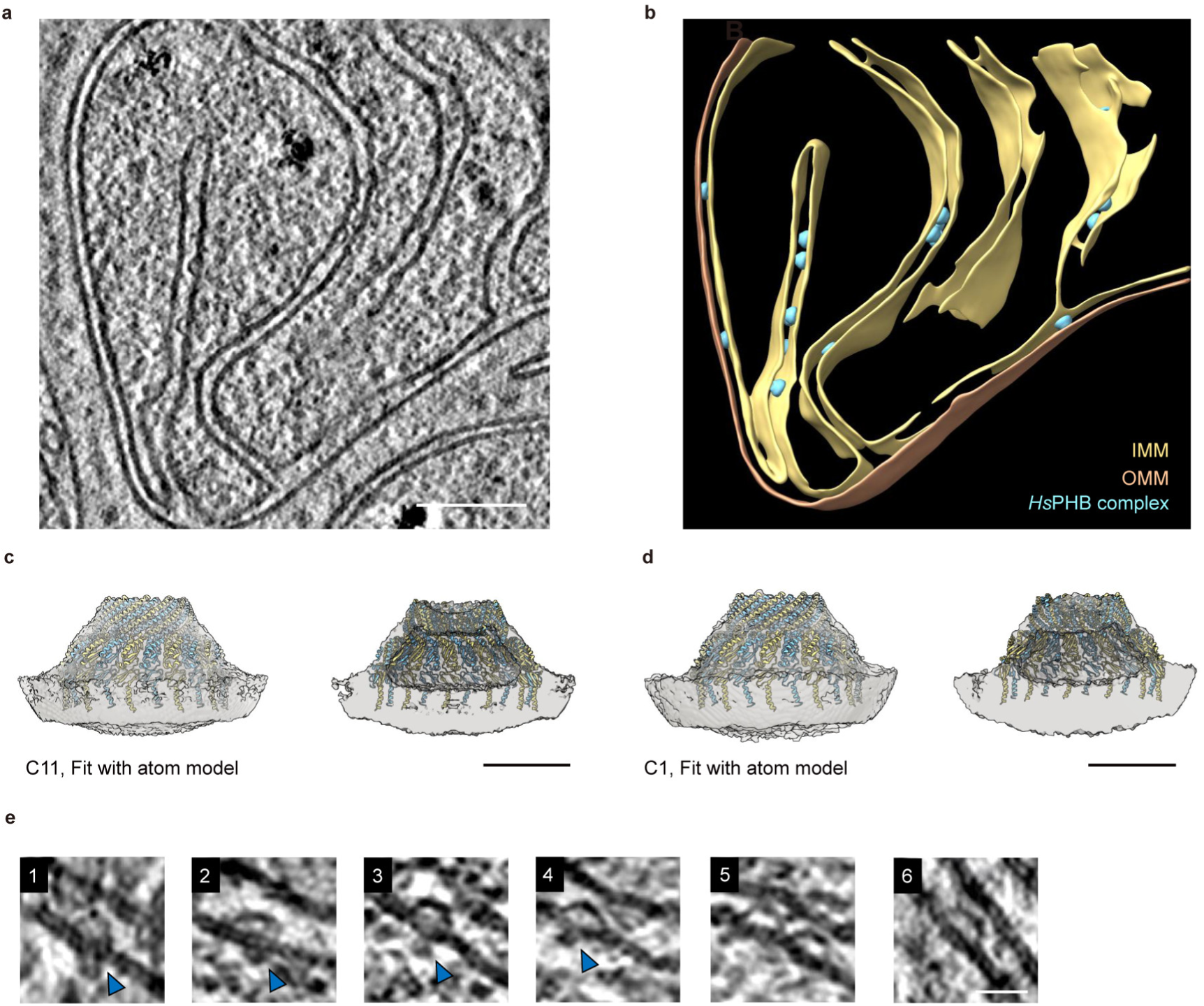
Cryo-ET reveals native architecture of *Hs*PHB complexes in human mitochondria. **a, b,** Tomographic analysis of mitochondrial membranes in HEK293T cells. Representative slice showing *Hs*PHB complexes decorating IMM (a) and corresponding 3D surface rendering (b). The *Hs*PHB complex, cyan; IMM, yellow; OMM, orange. Scale bar, 100 nm. **c, d,** Subtomogram averaging of *Hs*PHB complexes. Density maps refined with C11 (c) and C1 (d) symmetry, fitted with the atomic model. Scale bar, 10 nm. **e,** Gallery of the *Hs*PHB complex tomographic slices, showing matrix-facing densities (blue arrowheads) beneath in some cases. Scale bar, 20 nm.

Importantly, detailed examination of the tomographic data revealed characteristic densities in the mitochondrial matrix beneath the majorities of the PHB cages (Fig. 5e), consistent with nsEM observations of *m*-AAA proteases within PHB complexes. These cellular tomographic results confirm the structural organization of the PHB complex and support its potential role as an IMM spatial organizer, compartmentalizing functional modules such as m-AAA proteases.

## Discussion

Our study reveals the molecular mechanisms underlying protein homeostasis regulation in the inner mitochondrial membrane (IMM) through the PHB complex-mediated control of *m*-AAA protease activity. The IMM exhibits remarkably high protein density, with abundant respiratory chain complexes and other membrane proteins distributed all over the membrane surface. Such protein-enriched membrane environment may require sophisticated compartmentalization of membrane-embedded proteases to prevent uncontrolled proteolysis. Our structural and functional analyses demonstrate that the PHB complex assembles into a cage-like architecture that spatially restricts *m*-AAA protease activity, providing structural evidence for compartmentalized proteolysis in the IMM.

The high-resolution structure of the *Ct*PHB complex reveals a precisely organized cage-like architecture that creates defined membrane microdomains. Meanwhile, this cage-shaped assembly shares structural features with other SPFH family members, including HflK/C^31,32^ and flotillin-1/2 complexes^36,37^, pointing to a common structural framework in membrane microdomain formation. Given the high sequence homology among prohibitins from species from yeast all the way to mammals (Fig. S7), it is foreseeable that PHBs from other species should form an analogous cage of the similar size and subunit stoichiometry. Our cryo-ET data confirmed that *Hs*PHBs indeed forms similarly sized cages in the IMM of human mitochondria. While preparing this manuscript, using a subtomogram averaging approach, two research groups independently reported the *in situ* structural characterization of the *Hs*PHB complex. However, limited by the resolution achievable in their cellular cryo-ET data, one study has incorrectly suggested an oligomeric state of 11 subunit ^43^ for the *Hs*PHB complex. Another study proposed an oligomeric state of 24 subunits^44^, partially based on the prokaryotic HflK/C complex ^31,32^. Our *in vitro* data reveals the precise molecular organization comprising 22 subunits for the *Ct*PHB complex, which is likely true for other species as well. Moreover, previous studies have identified ∼120 kDa intermediates of newly imported *Sc*Phb1 and *Sc*Phb2 subunits in yeast, suggesting that the complex undergoes dynamic assembly and disassembly through regulated inter-subunit interactions^12^. Deletion of the CC domains prevents further assembly of the complex^12^, highlighting their essential role in complex formation. Our study further demonstrates that introducing charged mutations at the CC1-CC2 junction compromises complex integrity, suggesting that post-translational modification such as phosphorylation in this region could serve as a potential regulatory mechanism to modulate the assembly-disassembly cycle.

Consistent with the spatial regulatory model established for the HflK/C-FtsH system in bacteria^31,32^, our findings extend this conserved mechanism to the mitochondrial PHB-*m*-AAA protease supercomplex by providing direct structural evidence through nsEM analysis of *in vitro* purified supercomplexes and *in situ* cryo-ET visualization of the native IMM environment. Furthermore, our study identifies SIM as a critical element mediating the regulation of *m*-AAA protease by the PHB complex (Fig. 6a).

**Figure 6.**
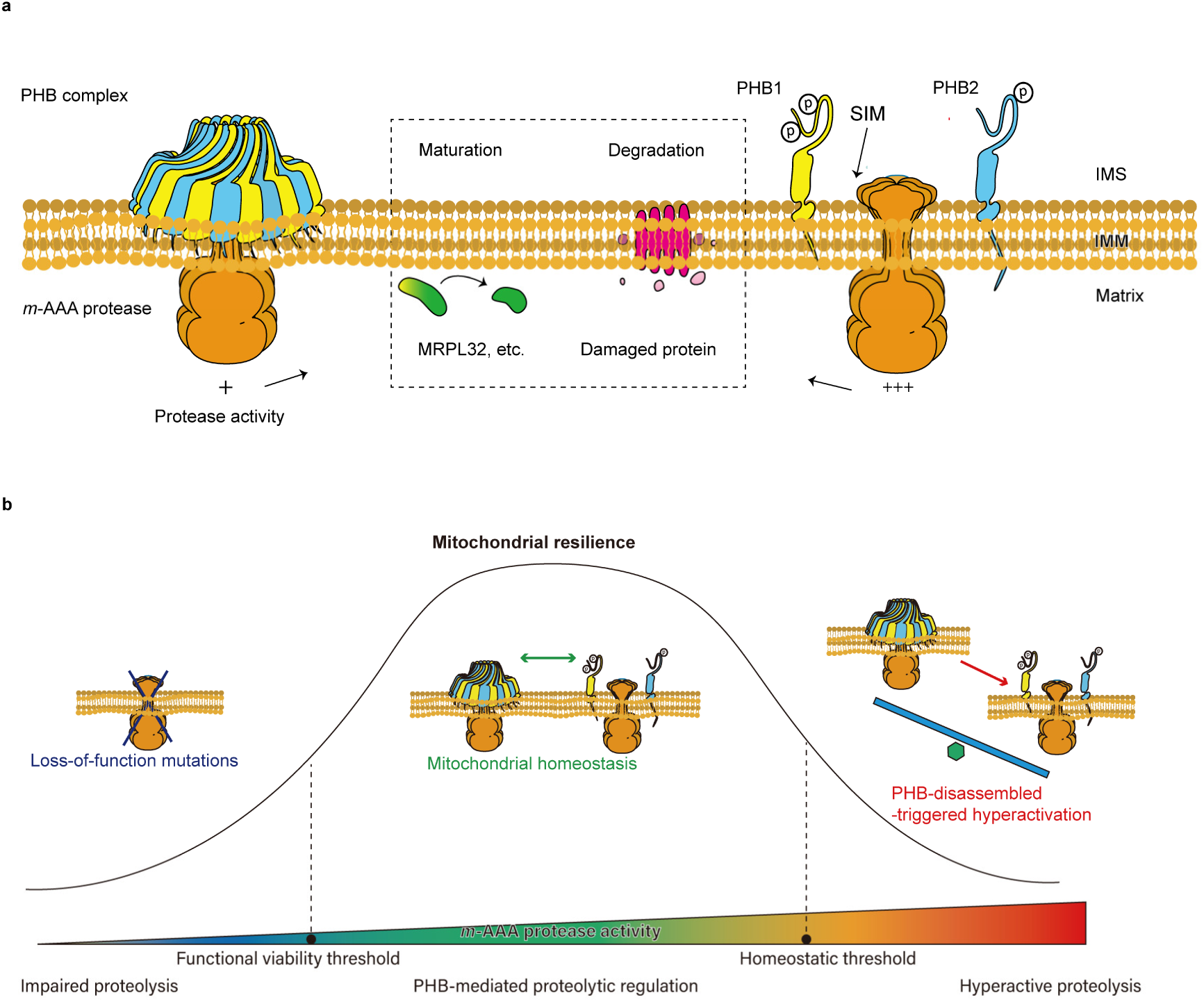
Model for PHB-mediated regulation of *m*-AAA protease activity in mitochondrial protein homeostasis. **a,** Schematic representation of PHB-mediated proteolytic regulation. Left: the PHB complex forms a cage-like structure that restricts *m*-AAA protease activity (+), enabling controlled substrate processing for protein maturation and degradation (boxed region). Right: SIM-mediated interaction between individual PHB subunits and *m*-AAA protease showing elevated protease activity (+++) when complex assembly is disrupted. **b,** Activity-dependent model of mitochondrial resilience. Proteolytic activity is required to be maintained between functional viability and homeostatic thresholds. Loss-of-function mutations cause impaired proteolysis (blue), while the PHB complex disassembly triggers hyperactive proteolysis (red). Balanced PHB-mediated regulation (green) maintains mitochondrial homeostasis through controlled proteolytic activity.

Previous studies established the essential role of *m*-AAA protease in maintaining mitochondrial function and homeostasis^8,23,45^. Guided by structural insights, our functional studies in yeast extend a critical requirement for properly balanced *m*-AAA protease activity in maintaining mitochondrial resilience (Fig. 6b). Specifically, reduced *m*-AAA protease activity (through IMSD deletion) leads to accumulation of misfolded and damaged proteins, while hyperactivation of *m*-AAA protease (through PHB1/2 deletion or SIM mutation) results in uncontrolled proteolysis and undesired degradation of functional proteins. Both scenarios ultimately compromise mitochondrial function and increase sensitivity to oxidative stress. In order to maintain homeostatic conditions, cells coordinate the expression of both PHB complex and *m*-AAA protease, as exemplified during diauxic shift when mitochondria undergo extensive proteome reorganization^29^. Notably, this regulatory mechanism may have important implications in cancer cells exposed to elevated ROS levels^46^. Indeed, increased PHBs expression has been reported in various cancer types^18–21^. Based on our model, the upregulation of PHBs might represent an adaptive response to maintain precise control over *m*-AAA protease activity under oxidative stress. Accordingly, targeting the PHB-*m*-AAA protease interaction emerges as a potential therapeutic strategy in cancer treatment, with the natural compound aurilide demonstrating initial proof-of-concept through its ability to disrupt this interaction and trigger apoptotic cell death^47^.

## Acknowledgements

We thank Dr. Chenkun Wang for advice on protein purification, Dr. Chengying Ma, and Dr. Ningning Li for their technical support in cryo-EM data processing, and all members of the Gao Lab and Guo Lab for stimulating scientific discussions and collaborative support. We acknowledge the Core Facilities at the School of Life Sciences, Peking University for assistance with negative-staining electron microscopy; the Cryo-EM Platform and the Electron Microscopy Laboratory of Peking University for assistance with cryo-EM data acquisition; the High-performance Computing Platform of Peking University for assistance with computation; and the National Center for Protein Sciences at Peking University for experimental support. This work was supported by National Natural Science Foundation of China (92354306 to N.G.), and by the Ministry of Science and Technology of China and Changping Laboratory. N.G. was supported in part by the Frontier Innovation Fund of Peking University Chengdu Academy for Advanced Interdisciplinary Biotechnologies.

## Author contributions

N.G. and Q.G. designed and supervised the project; D.L. carried out protein preparation, biochemical experiments and analyzed the Cryo-EM data with help from M.L. and L.Z.; P.W. and Z.C. performed cryo-ET experiments; D.L. and Z.C. prepared figures; D.L. prepared manuscript. N.G. built atomic model and analyzed structure with D.L.; N.G., Q.G. and M.L. revised the manuscript. All the authors approved the manuscript.

## Data and materials availability

The cryo-EM map and the structure coordinates have been deposited in the Electron Microscopy Data Bank and the Protein Data Bank under accession codes EMD-XXXX and XXXX.

## Supplementary Materials

Figs. S1 to S10

Table S1

## Methods

### Molecular cloning and generation of yeast mutant strains

All coding sequences used in this study were obtained either through gene synthesis or genomic DNA amplification. For proteins from *Chaetomium thermophilum* and *Homo sapiens*, genes encoding subunits of the PHB complex (*Ct*Phb1: G0SEE1; *Ct*Phb2: G0SCJ5; *Hs*PHB1: P35232; *Hs*PHB2: Q99623) and *m*-AAA proteases (*Ct*Rca1: G0S820; AFG3L2: Q9Y4W6) were synthesized by GenScript based on their amino acid sequences from UniProt database with codon optimization. The corresponding *Saccharomyces cerevisiae* genes were amplified from genomic DNA, and their protein sequences can be found in UniProt database (*Sc*Phb1: P40961; *Sc*Phb2: P50085; Yta12: P40341; Yta10: P39925; MrpL32: P25348).

For protein overexpression in HEK293F system, the gene encoding chimeric peptide containing *Hs*PHB1 (C-terminal Strep II tag) and *Hs*PHB2 (C-terminal FLAG-tag) connected by P2A self-cleaving peptide sequence (ATNFSLLKQAGDVEENPGP) was cloned into PKH3 vector under CMV promoter. The genes encoding phosphomimetic variants (*Hs*PHB1^Y259E^, *Hs*PHB1^Y265E^, *Hs*PHB2^Y248E^ and triple mutant *Hs*PHB^3E^) were generated in this backbone. The gene encoding engineered AFG3L2 (C-terminal Strep II tag; inactivating mutations E408Q for ATPase and E575Q for protease activities) was cloned into PKH3 backbone with CAG promoter, based on which two variants, AFG3L2^ΔSIM^ (deletion of residues 201–210) and AFG3L2^ΔIMSD^ (replacement of residues 164–250 with GGGGGS), were subsequently constructed.

For protein overexpression in yeast system (W303-1B background), the genes encoding the PHB complex components from different species were cloned into pESC vectors with GAL1/GAL10 bidirectional promoters: *Ct*Phb1, *Sc*Phb1 and *Hs*PHB1 (N-terminal FLAG-tag) under GAL1 promoter, while *Ct*Phb2, *Sc*Phb2 and *Hs*PHB2 (N-terminal Strep II-tag) under GAL10 promoter, based on which the gene encoding *Ct*Phb2^Y266E^ variant was constructed to examine charge effects on complex assembly. The genes encoding engineered *Ct*Rca1 and AFG3L2 (both with C-terminal Strep II-tag) were constructed by replacing their N-terminal disordered regions (residues 1–287 and 1– 139, respectively) with *Saccharomyces cerevisiae* Yta10 mitochondrial targeting sequence (residues 1–61). Catalytically inactive *Ct*Rca1 variant was generated by introducing E559Q/E725Q mutations, while AFG3L2 carried the aforementioned inactivating mutations. Based on these constructs, two additional *Ct*Rca1 variants were generated: *Ct*Rca1^ΔSIM^ (replacement of residues 344–354 with GGGGGS linker) and *Ct*Rca1^ΔIMSD^ (replacement of residues 311–400 with GGGGGS linker). For PHB-*m*-AAA protease supercomplex expression, the genes encoding *m*-AAA protease components and GEV (engineered fusion protein Gal4dbd.ER.VP16^48^) were introduced into PHB-expressing vectors as GAL1/GAL10 promoter-driven and TEF promoter-driven expression cassettes, respectively. GEV functions as a transcription factor that, upon allosteric activation by β-estradiol, binds to GAL promoters to induce target gene expression^48^.

For complementation studies and protease activity assays, the genes encoding *Sc*Phb1, *Sc*Phb2, yta12, and MrpL32 were introduced into pESC backbone under TEF promoter control, respectively.

Deletion strains (W303-1B background) lacking *Sc*Phb1, *Sc*Phb2, or Yta12 (*Δscphb1*, *Δscphb2*, *Δyta12*), and domain-specific mutants *yta12^ΔIMSD^*(residues 199–290) and *yta12^ΔSIM^* (residues 236–251) replaced with GGGGGS linkers were generated using established yeast genome engineering system based on CRISPR-Cas9^49^ and confirmed by sequencing.

### Cell Culture and protein expression

HEK293F cells were cultured in FreeStyle 293 expression medium (Thermo Fisher) under standard conditions (37°C, 5% CO_2_, 125 rpm). Cells were maintained in exponential growth phase by passaging at 1.0–1.5 × 10^6^ cells/ml. For protein expression, cultures at 1.5–2.0 × 10^6^ cells/ml were transfected using polyethylenimine (PEI, Polysciences) at a DNA: PEI mass ratio of 1:3. For co-transfections, plasmids were combined at equimolar ratios (1 mg total DNA per liter of culture). DNA-PEI mixtures were incubated in fresh medium for 30 min at room temperature before adding to cultures. Cells were harvested 48 h after transfection.

Expression strains (W303-1B) were transformed with plasmids encoding PHB complexes (*Ct*PHB, *Hs*PHB and *Sc*PHB), *m*-AAA proteases (*Ct*Rca1 and AFG3L2), or PHB-*m*-AAA protease supercomplexes (*Ct*PHB-*Ct*Rca1 and *Hs*PHB-AFG3L2). Transformants were cultured overnight in SC-his (for PHB complexes and PHB-*m*-AAA protease supercomplexes) or SC-ura (for *m*-AAA proteases) media. For individual expression of the PHB complexes or *m*-AAA proteases, overnight cultures were diluted 1:10 in YPGal medium containing 2% (w/v) galactose and grown for 18h. For PHB-*m*-AAA protease supercomplexes, cells were first diluted 1:10 in YPEG medium and grown for 10 h, followed by induction with 100 nM β-estradiol for 10–12 h.

### Mitochondrial Isolation

For mitochondrial isolation from HEK293F cells, harvested cells were processed by using Dounce homogenization (50 strokes) in buffer A (250 mM Sucrose, 50 mM HEPES-KOH, pH 7.4) containing 1% (v/v) protease inhibitors cocktail (Mei5bio). Mitochondrial fractions were pelleted by centrifugation (15,000× g, 30 min; 25.5A rotor, Beckman Coulter) resuspended in buffer B (100 mM KOAc, 10 mM Mg(OAc)_2_, 25 mM HEPES-KOH, 10% glycerol, pH 7.8), and stored at −80°C.

For mitochondrial isolation from yeast strains, harvested cells were suspended in buffer C (600 mM Sorbitol, 100 mM HEPES-KOH, pH 7.4) containing 1% (v/v) protease inhibitors cocktail (Mei5bio) and 0.2% (w/v) BSA (Mei5bio). The samples were cryo-milled (6875 Freezer/Mill), thawed on ice, and subjected to centrifugation (3,000× *g*, 30 min; Eppendorf 5804R) to remove cellular debris. The supernatant was further centrifuged (20,000× *g*, 60 min; 25.5A rotor, Beckman Coulter) to obtain crude mitochondria. Purified mitochondria were isolated by density gradient centrifugation through a discontinuous sucrose gradient^50^. The purified mitochondrial fraction was collected at the 32%/60% interface, resuspended in buffer B, and stored at −80°C.

### Protein purification

All protein complexes, including PHB complexes (*Sc*PHB, *Ct*PHB, *Hs*PHB), *m*-AAA proteases (*Ct*Rca1 and AFG3L2), and their supercomplexes (*Ct*PHB-*Ct*Rca1 and *Hs*PHB-AFG3L2), were expressed in *Saccharomyces cerevisiae* and purified at 4℃ using distinct protocols.

For PHB complexes or *m*-AAA proteases, proteins were extracted from whole cell membrane of harvested cells resuspended in buffer C. Cells were disrupted using a high-pressure homogenizer (JNBIO) at 1,800 bar with two passages. The homogenate was cleared by centrifugation (3,000× *g*, 10 min; Eppendorf 5804R), followed by membrane fraction collection through ultracentrifugation (150,000× *g*, 60 min; 70Ti rotor, Beckman Coulter). For PHB-*m*-AAA protease supercomplexes, mitochondrial membrane fractions were isolated following the mitochondrial isolation protocol described above.

Membrane proteins were extracted using complex-specific detergent conditions: *Ct*PHB/*Sc*PHB with 1% (w/v) DDM (Anatrace), AFG3L2/*Ct*Rca1 with 1% (w/v) LMNG (Anatrace), *Hs*PHB with 3% (w/v) OG (Anatrace) and *Ct*PHB-Rca1/*Hs*PHB-AFG3L2 supercomplexes with 1% (w/v) digitonin (Biosynth). Membrane pellets were resuspended in buffer B at 1:5 (v/v) ratio (1:10 for digitonin samples). Solubilization was performed with magnetic stirring for 2 h, followed by ultracentrifugation (150,000× *g*, 35 min; 70Ti rotor, Beckman Coulter) to remove insoluble material.

For purification, solubilized proteins were incubated for 2 h with pre-equilibrated Anti-Flag M2 Affinity Gel (Sigma-Aldrich) for PHB complex or Streptactin Beads 4FF (Smart-Lifesciences) for *m*-AAA proteases. After washing with buffer B containing 2× critical micelle concentration of the same detergent used for solubilization, proteins were eluted in wash buffer supplemented with 0.2 mg/ml 3×Flag peptide (NJPeptide) for the PHB complexes or 20 mM d-Desthiobiotin (Sigma-Aldrich) for *m*-AAA proteases. For PHB-*m*-AAA protease supercomplexes, a two-step affinity purification strategy was employed. Samples were first purified using Anti-Flag M2 Affinity Gel to capture Flag-tagged PHB complexes, followed by Streptactin Beads 4FF purification to isolate Strep-tagged *m*-AAA proteases and their associated PHB complexes.

Purified proteins were concentrated using a 100 kDa MWCO spin concentrator (Millipore) and further fractionated by 20–50% glycerol density gradient ultracentrifugation in buffer B (100,000× *g*, 14 h; TLS-55 rotor, Beckman Coulter) to enhance sample homogeneity.

### Pull-down assay

Physical interactions between PHB complexes and *m*-AAA protease variants (wild-type, ΔSIM and ΔIMSD) were analyzed by pull-down assays. *Ct*PHB-*Ct*Rca1 and *Hs*PHB-AFG3L2 complexes were expressed in yeast and HEK293F, respectively, as described above. Mitochondrial membrane fractions were solubilized in buffer B containing 1% (w/v) digitonin at 4°C. Protein concentrations were determined using NanoDrop Lite spectrophotometer (Thermo Scientific) and normalized with buffer B containing 0.1% (w/v) digitonin. Equal amounts of solubilized proteins were incubated with Anti-Flag M2 Affinity Gel for 2 h at 4°C. After washing with buffer B containing 0.1% (w/v) digitonin, bound proteins were resuspended in the same buffer, mixed with 5× SDS-PAGE sample buffer, and heated (95°C, 10 min). Samples were centrifuged (12,000× *g*, 5 min; Eppendorf 5424R) and analyzed by SDS-PAGE followed by immunoblotting.

### The PHB complex assembly states assay

The oligomeric states of PHB complex variants containing CC-region charge mutations (*Ct*Phb2^Y266E^, *Hs*PHB1^Y259E^, *Hs*PHB1S^265E^, *Hs*PHB2^Y248E^ and *Hs*PHB2^Y248E^) were analyzed by glycerol density gradient centrifugation. The *Ct*Phb2^Y266E^ variant was expressed in *Saccharomyces cerevisiae*, while *Hs*PHB variants were expressed in HEK293F cells. All procedures were performed at 4°C. Mitochondrial membrane fractions were solubilized in buffer B containing 1% (w/v) digitonin for 1 h. Following ultracentrifugation (160,000× *g*, 15 min; TLA55 rotor, Beckman Coulter), the supernatant was fractionated on a 20–50% glycerol density gradient in buffer B containing 0.1% (w/v) digitonin (100,000× *g*, 14 h; TLS55 rotor, Beckman Coulter). Gradient fractions were analyzed by SDS-PAGE and immunoblotting to evaluate the PHB complex assembly states.

### MrpL32 processing assay

To assess *m*-AAA protease activity, MrpL32 processing was monitored in various yeast strains (wild-type, *Δscphb1*, *yta12^ΔIMSD^*, and *yta12^ΔSIM^*) transformed with pESC-HIS-TEF-MrpL32^CStrep^. All procedures were performed at 4°C unless specified otherwise. Yeast cultures were grown in YPG medium (starting OD_600_ ∼0.1) for 14 h at 30°C. For small-scale mitochondrial isolation, cells were harvested and resuspended in buffer C containing 1% (v/v) protease inhibitors. Cells were disrupted using glass beads through 10 cycles of alternating vortexing (15 s) and ice incubation (15 s). After removing cell debris (3,000*×g*, 10 min; Eppendorf 5804R), crude mitochondria were isolated by centrifugation (16,000× *g*, 30 min; Eppendorf 5804R) and resuspended in buffer B. Samples were mixed with 5× SDS-PAGE sample buffer, heated (95°C, 10 min), and centrifuged (12,000× *g*, 5 min; Eppendorf 5424R). MrpL32 precursor and mature forms were analyzed by SDS-PAGE and immunoblotting. Band intensities were quantified using ImageJ software, with precursor-to-total protein ratios calculated from four independent biological replicates. Statistical significance was assessed using the one-way ANOVA test.

### Yeast oxidative stress sensitivity assay

Yeast strains (wild-type, *Δscphb1*, *Δscphb2*, *yta12^ΔIMSD^*, *yta12^ΔSIM^*, *Δyta12* and their complemented derivatives) were evaluated for oxidative stress sensitivity. Cells were grown overnight at 30°C in YPD or SC-HIS media (selective medium for complementation plasmids with HIS3 marker), diluted to OD_600_ = 0.1 in fresh YPD medium, and cultured at 30°C for additional 14 h. Cell suspensions were harvested, washed with sterile phosphate-buffered saline (PBS, Thermo Fisher Scientific), and normalized to equal cell densities based on OD_600_. Serial dilutions (10^0^ to 10^-5^) were prepared in PBS, and 3 μl of each dilution was spotted onto YPD or YPG plates (with or without 2 mM H_2_O_2_). Colony growth was monitored after 48h (YPD) or 72 h (YPG) incubation at 30°C.

### Electron microscopy sample preparation

For nsEM sample preparation, protein samples (4 μl, ∼0.1 mg/ml) were applied to glow-discharged copper grids (Emcn) and stained with 2% uranyl acetate. Single-frame micrographs were collected on an FEI Tecnai T20 TEM operated at 120 kV, with a magnification of 62,000× corresponding to a physical pixel size of 1.57 Å/pixel. A total of 200 micrographs were recorded for subsequent analysis.

For Cryo-EM sample preparation, the *Ct*PHB complex samples (4 μl, ∼5 mg/ml) were applied to gold grids (Quantifoil, R1.2/1.3 300-mesh) and vitrified using an FEI Vitrobot Mark IV (8°C, 100% humidity) by plunge-freezing in liquid ethane. Grids were initially screened on an FEI Talos Arctica microscope equipped with an FEI Ceta camera at 200 kV. Data collection was performed on an FEI Titan Krios microscope operated at 300 kV, equipped with a Gatan K2 summit direct electron detector and GIF energy filter. The movies were acquired using SerialEM^51^ at a dose rate of 6.55 e−/s/Å^2^ and an exposure time of 8s with 32 frames. The data was collected at the magnification of 130,000× corresponding to a physical pixel size of 1.052 Å/pixel and with defocus ranging from −1.2 to −1.8 μm. 2900 of raw movies were collected in total.

### Electron microscopy data processing

For nsEM data, initial particles were manually picked from raw micrographs using RELION 4.0^52^ to generate templates for automated particle picking. Auto-picked particles underwent reference-free 2D classification to characterize the structural organization of the complex.

For cryo-EM data, movie stacks were motion-corrected and dose-weighted using MotionCor2^53^. Contrast transfer function (CTF) parameters were estimated using Gctf^54^. All subsequent processing steps were performed using RELION 4.0. Initial 2D references were generated from manually picked particles for automated particle picking. Auto-picked particles underwent multiple rounds of reference-free 2D classification. An initial 3D model was generated from selected particles. Particles from well-defined classes were subjected to initial 3D classification. Classes showing intact structures proceeded to additional rounds of 3D classification or refinement. The final 3D refinement with global mask included 64,723 particles, yielding a 7.2 Å resolution map. Subsequent CTF refinement, Bayesian polishing and post-processing improved the resolution to 4.5 Å. To enhance the *Ct*PHB complex density, Hsp60 density was subtracted followed by masked 3D classification. The final refinement using 10,623 particles with C11 symmetry achieved 3.2 Å resolution based on the gold-standard FSC = 0.143 criterion. Local resolution distribution was estimated using the deep learning-based CryoRes algorithm^55^.

### Atomic model building and refinement

Initial model building of the C11-symmetric *Ct*PHB complex began with de novo modeling of *Ct*Phb1 and *Ct*Phb2 subunits using Coot software^56^. Additional copies of subunits were placed in the density map through rigid-body fitting, followed by manual adjustments based on local density features to generate the complete 22-mer atomic model. The model underwent multiple rounds of refinement using Phenix.real_space_refinement in the Phenix software suite^57^, alternating with manual adjustments to optimize fit to the electron density map while maintaining ideal geometric restraints. Model quality was assessed using MolProbity validation tools^58^.

### *In situ* cryo-ET analysis of *Hs*PHB complexes

HEK293T cells were seeded into new wells of 6-well plates containing glow-discharged grids (Gold mesh micro-gate film, T10012A; Beijing XXBR Technology Co., Ltd.) at a density of 3 × 10⁵ cells/well. They were then allowed to attach for 20 h in culture before vitrification. For vitrification, the grids were blotted from the back-side by replacing the front-side filter paper with parafilm. Post-blottling, the grids were immediately plunged into liquid ethane using a Vitrobot Mark IV (Thermo Fisher).

The grids were loaded into a sample shuttle and transferred into the cryo-focused ion beam/scanning electron microscope (cryo-FIB/SEM; Aquilos 2, Thermo Fisher) for thinning as described^59^. In brief, platinum layers were deposited via sputter coating and gas injection to reduce charging effects. During milling, the ion current was decreased stepwise, with a final current of 50 pA used for fine milling. Resulting lamellae had a mean thickness of 100–300 nm, suitable for subsequent tomographic data collection.

Grids were loaded into the Titan Krios G3 TEM equipped with a Gatan post-column energy filter and K3 direct electron detector. Tilt series were collected using SerialEM^51^ between −60° and +36° with an increment of 3 degrees, employing a dose-symmetry scheme^60^. This resulted in a total dose of 110 e⁻/Å² per tilt series. The pixel spacing was set at 1.37 Å, and the defocus was between −5 μm and −7 μm.

Raw data were pre-processed using TOMOMAN^61^ before being imported into IMOD software^62^ for alignment and reconstruction. Tilt series alignment utilized a fiducial-less patch-tracking mode, followed by full tomogram reconstruction through weighted back projection.

In total, 450 dome-like particles were manually selected from 17 mitochondria-containing tomograms. For structural analysis, subtomograms were extracted using Warp^63^, the resulting bin two subtomograms were subsequently classified and averaged using RELION^64^. The initial classification was performed without imposing any symmetry, resulting in a dome-like structure that matched the overall shape of the prohibitin complex. C11 symmetry was then applied for final refinement.

For visualization, mitochondria were segmented with MemBrain^65^ and *Hs*PHB complexes were positioned into the tomograms using orientation and positional information derived from subtomogram averaging. ChimeraX was used for final 3D rendering^66^.

