## Supplementary Materials for "Structural basis for prohibitin-mediated regulation of mitochondrial *m*-AAA protease"


#### This PDF file includes:

Figs. S1 to S10

Table S1

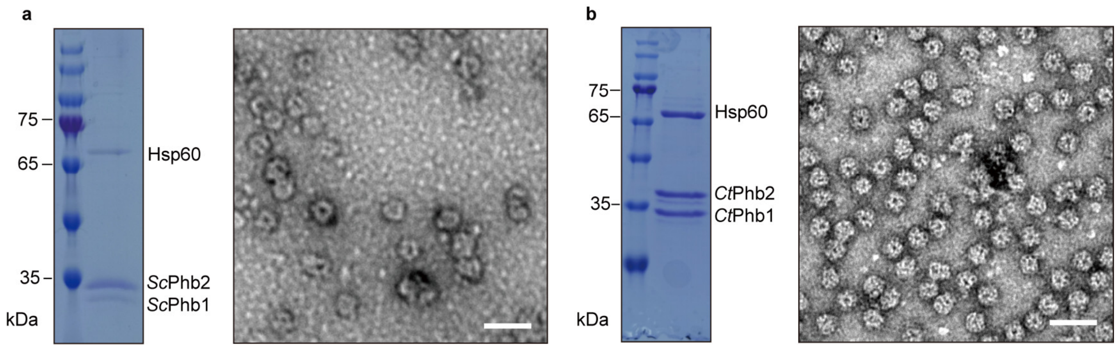


#### Figure S1. Sample quality assessment of the *Sc*PHB and *Ct*PHB complex expressed in *Saccharomyces cerevisiae*

**a, b,** SDS-PAGE (left) and nsEM analysis (right) of purified *Sc*PHB (a) and *Ct*PHB (b) complexes. SDS-PAGE shows co-purified Hsp60 (65 kDa) with both *Sc*PHB1/2 and *Ct*PHB1/2 (35 kDa). Scale bars, 50 nm.

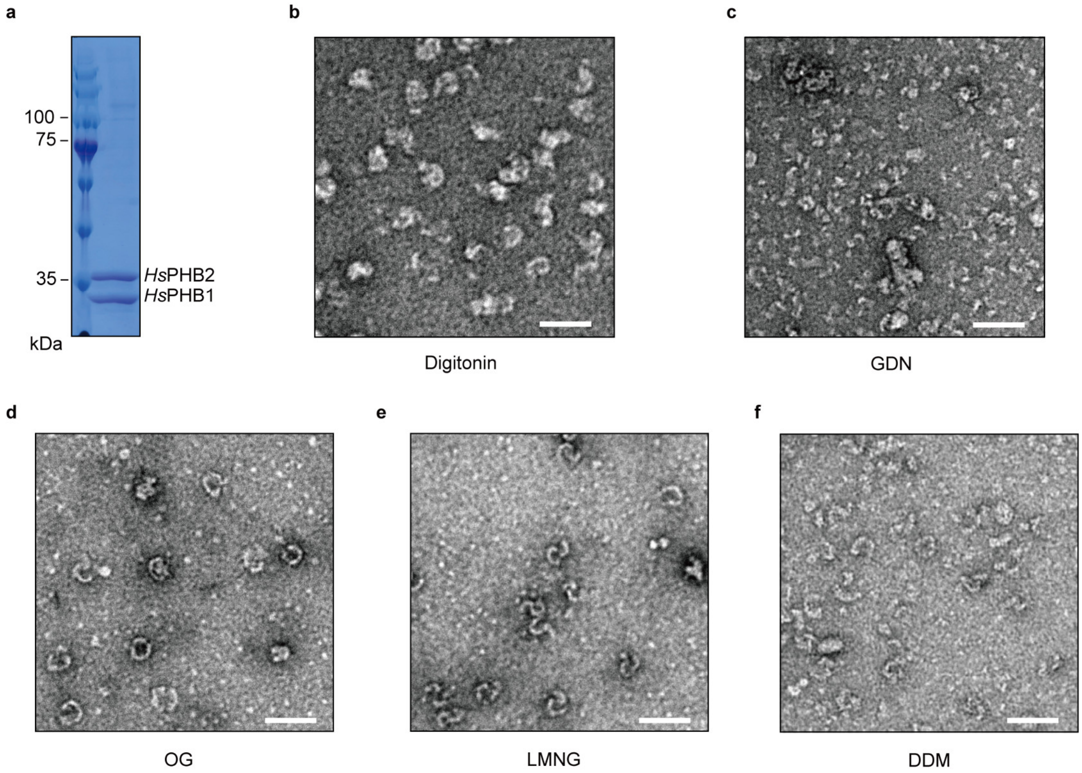


#### Figure S2. Sample homogeneity analysis of the *Hs*PHB complex in different detergents

**a,** SDS-PAGE analysis of purified *Hs*PHB complexes.

**b–f,** nsEM analysis of *Hs*PHB complexes in various detergents: digitonin (b), GDN (c), OG (d), LMNG (e), and DDM (f). Scale bars, 50 nm.

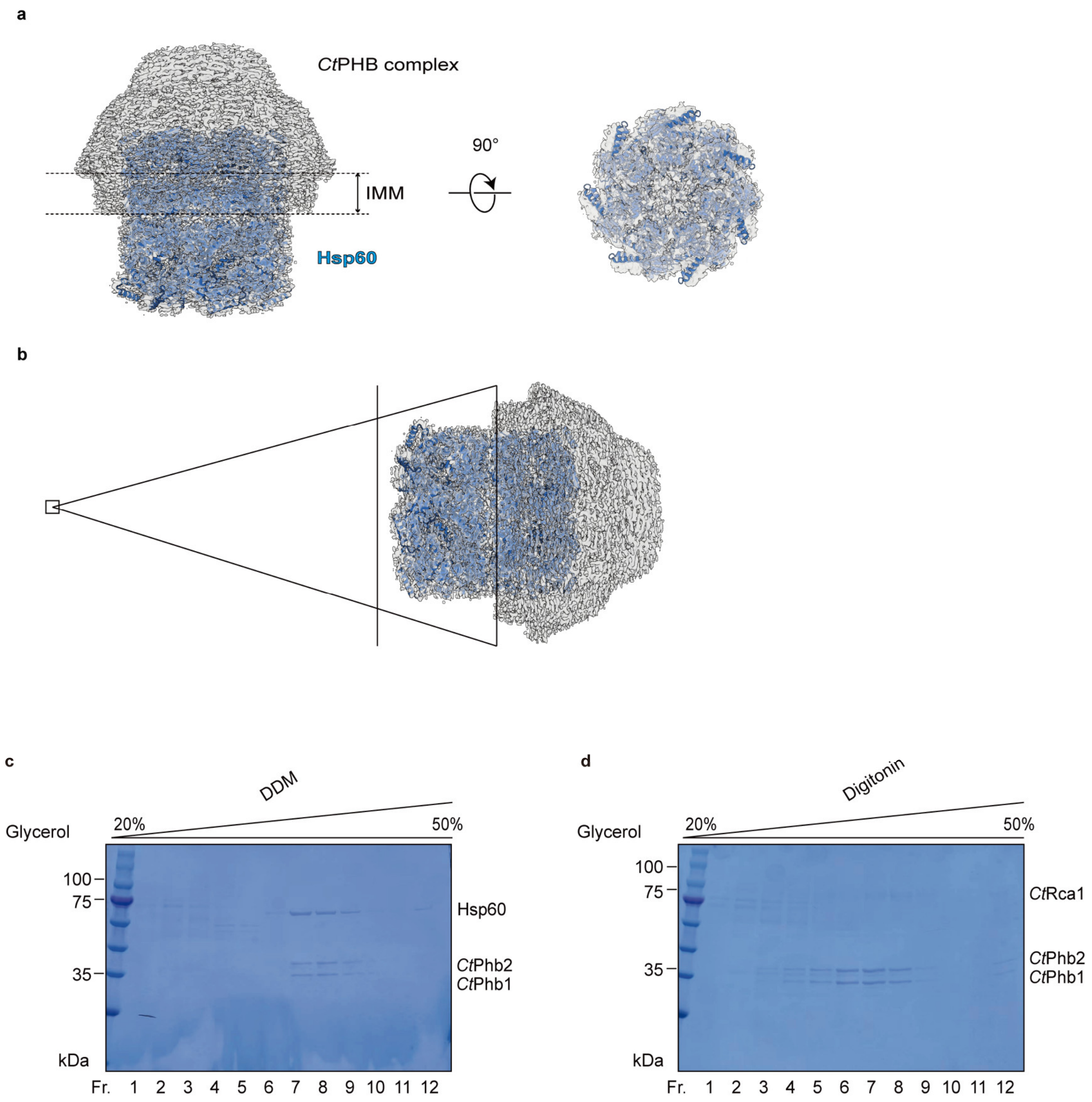


### Figure S3. Non-physiological association between Hsp60 and the *Ct*PHB complex

**a,** Side and bottom views (rotated 90°) of cryo-EM density map (grey) with docked AlphaFold-predicted Hsp60 double-ring tetradecamer model (blue). IMM position is indicated.

**b,** Schematic illustration showing the projection diagram and cross-sectional plane (vertical line) used to obtain the bottom view in a.

**c, d,** Glycerol gradient centrifugation (20–50%) analysis of *Ct*PHB-*Ct*Rca1 supercomplexes purified in DDM (c) or digitonin (d). DDM solubilizes the complex's internal lipids, allowing non-specific Hsp60 insertion, while mild detergent digitonin helps maintain both *Ct*PHB-lipid association and *Ct*PHB-*Ct*Rca1 interaction (Fr., fraction).

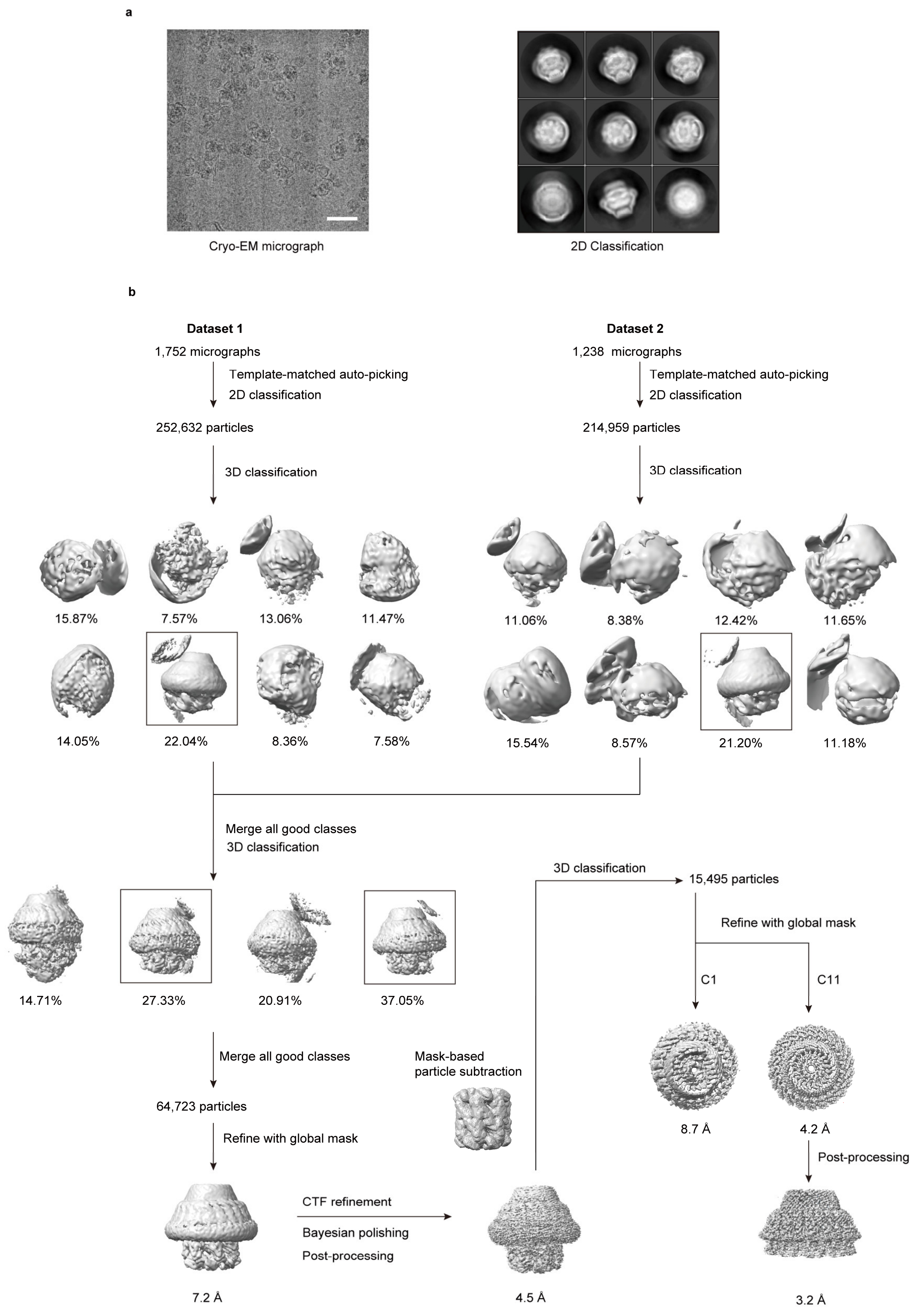


### Figure S4. Cryo-EM data processing workflow for the *Ct*PHB complex

**a,** Representative cryo-EM micrograph (left) and 2D class averages (right) of the *Ct*PHB complex. Scale bar, 50 nm.

**b,** Processing scheme for two independent datasets (Dataset 1: 1,752 micrographs; Dataset 2: 1,238 micrographs). Initial particle picking and 3D classification yielded selected classes (black boxes) that were combined for further processing. Sequential refinement steps improved the resolution from 7.2 Å to 4.5 Å. The final reconstruction after Hsp60 density subtraction achieved 3.2 Å. Percentage values indicate particle distribution in 3D classes.

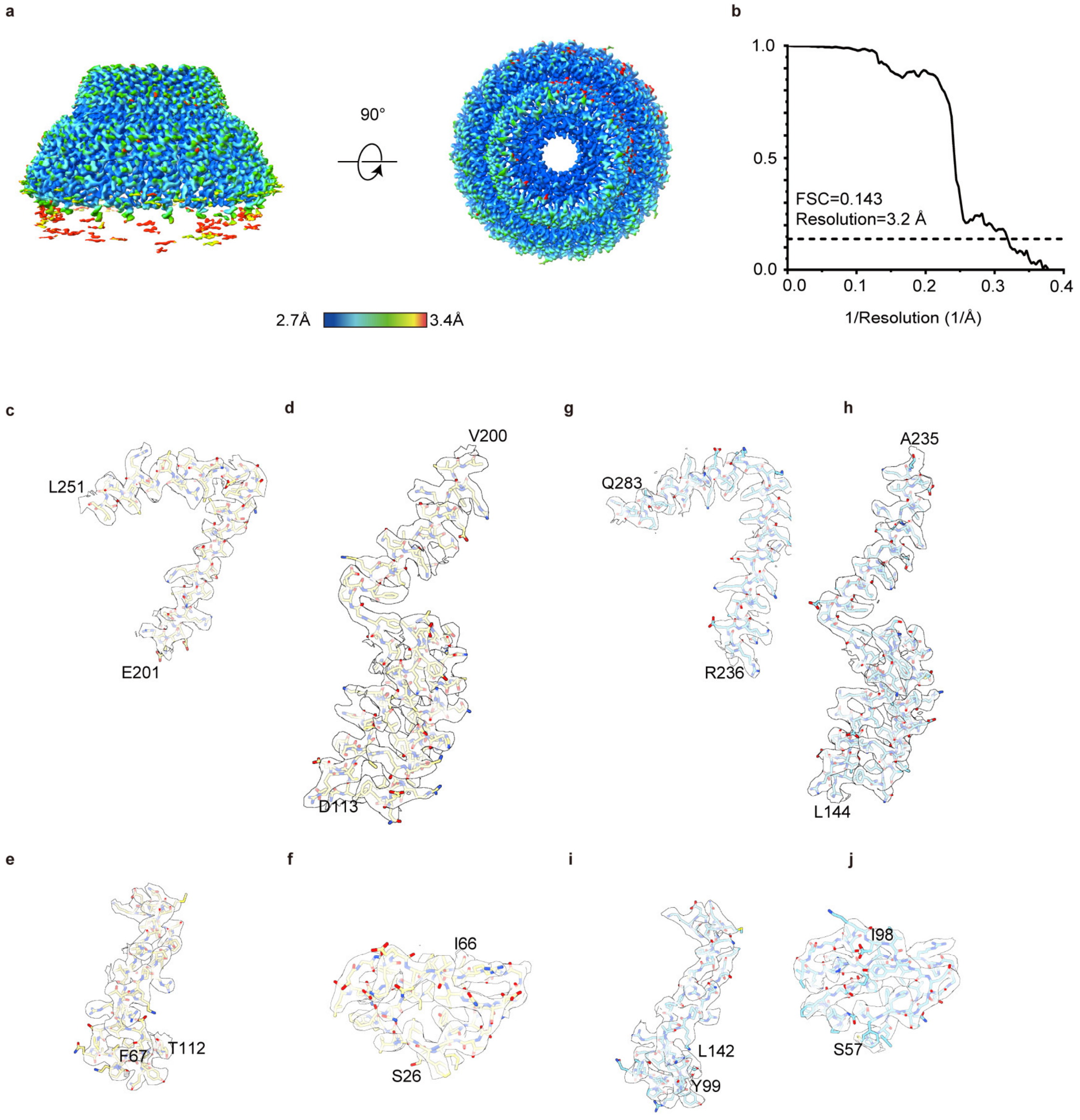


#### Figure S5. Cryo-EM Density Map Quality Assessment of the *Ct*PHB Complex

**a,** Local resolution distribution mapped onto the density in side and top views (90° rotation). The color scale ranges from 2.7 Å (blue) to 3.4 Å (red).

**b,** Gold-standard FSC curve showing overall resolution of 3.2 Å at FSC = 0.143.

**c–f,** Representative cryo-EM density maps (mesh) with fitted atomic models for *Ct*Phb1 segments: E201–L251 (c), D113–V200 (d), F67–T112 (e), and S26–I66 (f).

**g–j,** Representative density maps for *Ct*Phb2 segments: R236–Q283 (g), L144A235 (h), Y99–L142 (i), and S57–I98 (j).

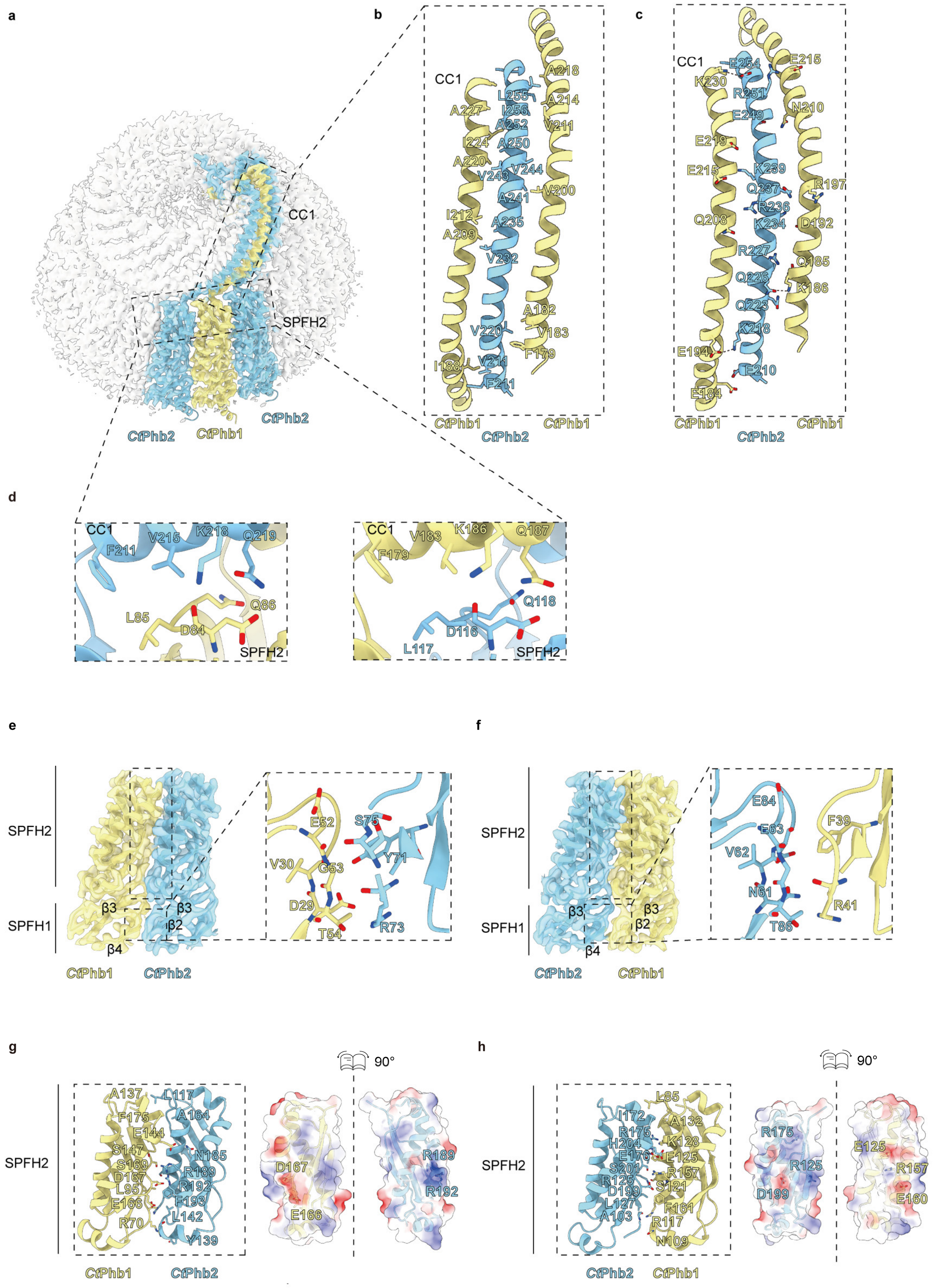

#### Figure S6. Structural analysis of interfaces between *Ct*Phb1 and *Ct*Phb2 subunits

**a,** Structure of a *Ct*Phb2-*Ct*Phb1-*Ct*Phb2 segment (*Ct*Phb2 in cyan, *Ct*Phb1 in yellow) within the complex (white), defining the P2-P1 and P1-P2 interfaces.

**b, c,** Helical interactions in the CC1 domain between *Ct*Phb1 (yellow) and *Ct*Phb2 (cyan), showing hydrophobic contacts between α-helices (b); polar interactions with hydrogen bonds indicated by dashed lines (c). Key interacting residues are shown as sticks

**d,** Close-up views of the CC1-SPFH2 connecting regions. At the P2-P1 interface (left), polar network formed by residues D84 and Q86 (*Ct*Phb1) with K218 and Q219 (*Ct*Phb2), and hydrophobic interactions formed by L85 (*Ct*Phb1) with F211 and V215 (*Ct*Phb2). At the P2-P1 interface (right), polar network formed by residues D116 and Q118 (*Ct*Phb2) with K186 and Q187 (*Ct*Phb1), and hydrophobic interactions formed by F179 and V183 (*Ct*Phb1) with L117 (*Ct*Phb2). Key residues are shown as sticks.

**e, f,** Interface analysis of SPFH1 domain showing interactions between β2-β3 and β3-β4 loops of adjacent subunits at P1-P2 (e) and P2-P1 (f) interfaces. Left panels show overall loop interactions, with zoomed-in views of key interfacial residues (shown as sticks) on the right.

**g, h,** Interface analysis of SPFH2 domain showing interacting interfaces in cartoon representation with key residues highlighted as stick models (left), and electrostatic surface potential analysis of the interfaces shown in 90° rotated "open-book" views (right) for P1-P2 (g) and P2-P1 (h) interfaces. Surface potentials are colored from negative (red) to positive (blue).

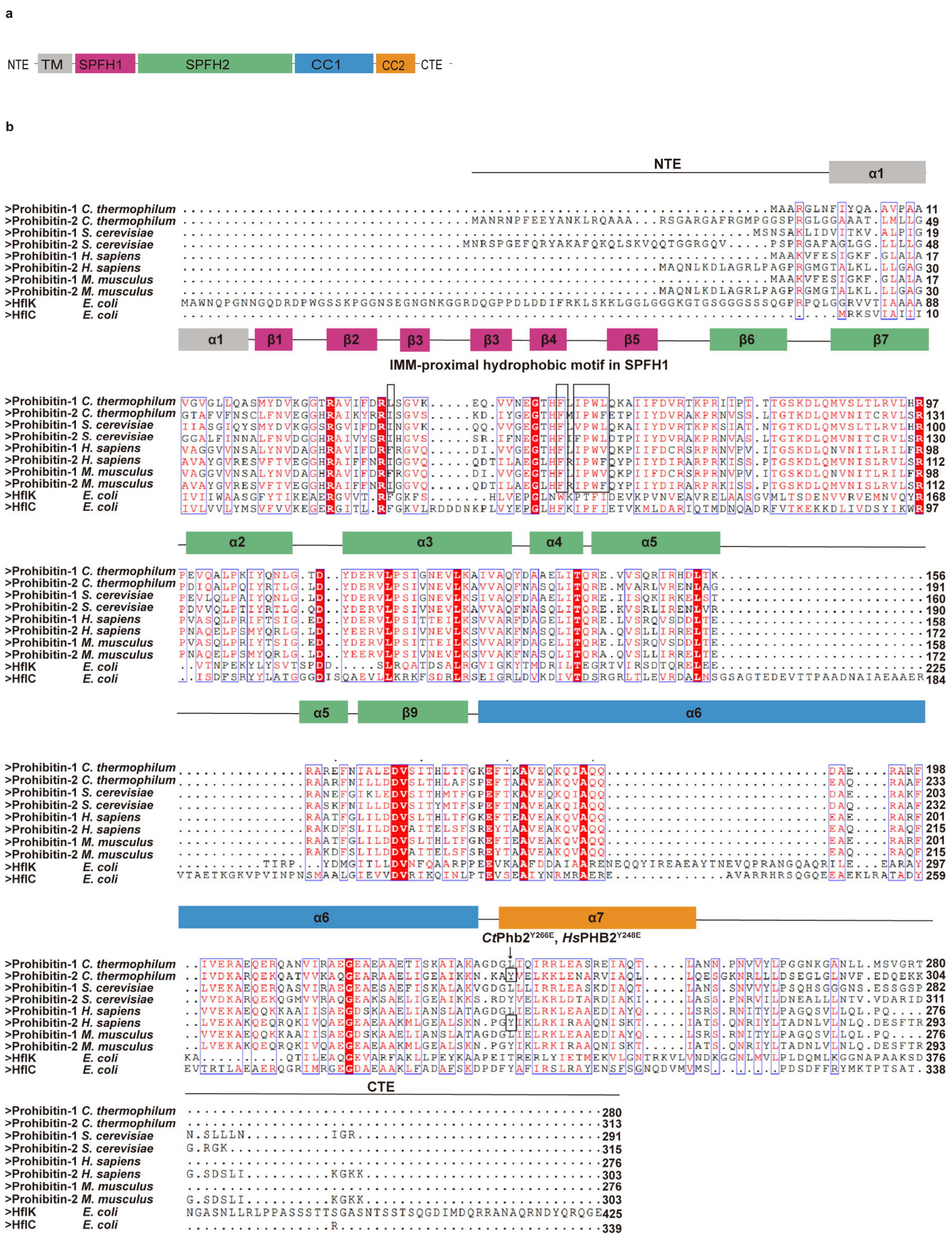


### Figure S7. Sequence conservation analysis of PHB proteins across species

**a,** Domain organization of PHB proteins showing N-terminal extension (NTE), transmembrane domain (TM), SPFH1 domain (pink), SPFH2 domain (green), coiled-coil domains CC1 (blue) and CC2 (orange) and C-terminal extension (CTE).

**b,** Multiple sequence alignment of PHB proteins from different species. Secondary structure elements are shown above the sequences and colored according to panel a. The IMM-proximal hydrophobic motif in SPFH1 domain and conserved corner residues in the CC domain (including *Ct*Phb2Y266 and *Hs*PHB2Y248) critical for complex integrity are highlighted.

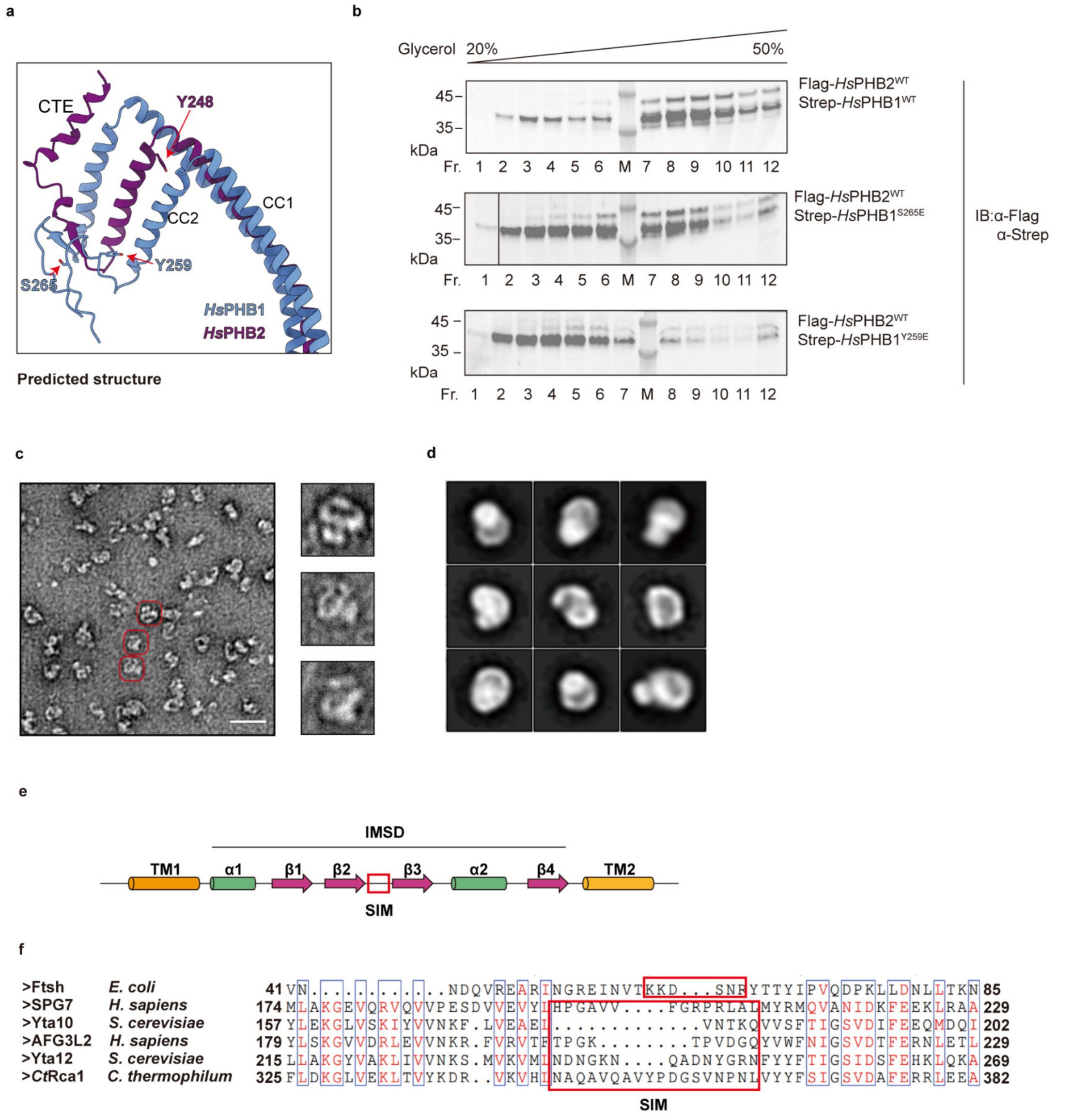


#### Figure S8. Structural analysis of *Hs*PHB complex assembly in CC region and its interaction with *m*-AAA protease

**a,** AlphaFold-predicted structure showing the CC domain corner region of *Hs*PHB complex. Key regulatory residues Y248 and Y259 are highlighted.

**b,** Glycerol gradient (20–50%) analysis of wild-type and mutant *Hs*PHB complexes. Flag-*Hs*PHB2$^{WT}$/Strep-*Hs*PHB1$^{WT}$ (top), Flag-*Hs*PHB2$^{WT}$/Strep-*Hs*PHB1$^{Y259E}$ (middle), and Flag-*Hs*PHB2$^{WT}$/Strep-*Hs*PHB1$^{S265E}$ (bottom) were analyzed by immunoblotting with anti-Flag and anti-Strep antibodies (Fr., fraction; M, marker).

**c–d,** nsEM analysis (c) and 2D class averages (d) of the *Hs*PHB-AFG3L2 complex. Scale bar, 50 nm.

**e,** Domain organization of the IMSD region showing transmembrane domains (TM1, TM2), α-helices (α1, α2), β-strands (β1–β4) and the SPFH-interacting motif (SIM).

**f,** Multiple sequence alignment of SIMs across species. Conserved residues are highlighted, and SIMs are boxed in red.

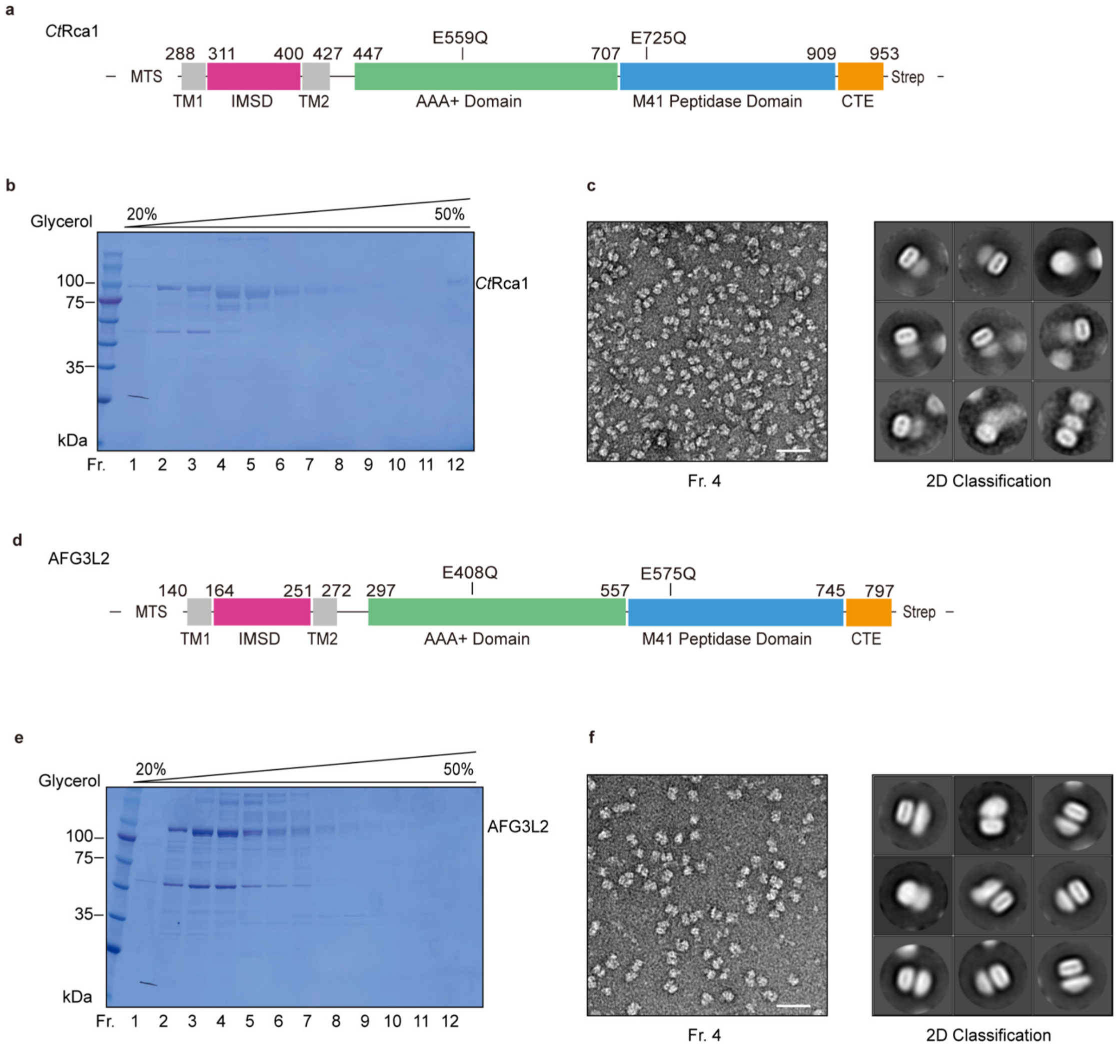


#### Figure S9. Expression and structural validation of *Ct*Rca1 and AFG3L2

**a,** Schematic representation of *Ct*Rca1 domain organization showing mitochondrial targeting sequence (MTS, corresponding to Yta12 residues 1–61), transmembrane domains (TM1, TM2), intermembrane space domain (IMSD), AAA+ domain with Walker B mutation (E559Q), M41 peptidase domain with active site mutation (E725Q) and C-terminal extension (CTE).

**b, c,** Sample quality assessment of purified *Ct*Rca1. Glycerol gradient (20–50%) analysis showing protein distribution (b). Representative nsEM micrograph (left) and 2D class averages (right) of fraction 4 (Fr. 4), revealing characteristic double-layered architecture (c). Scale bar, 50 nm

**d, f,** Parallel analysis of AFG3L2. Domain organization with corresponding mutations E408Q and E575Q (d). Glycerol gradient analysis (e). NsEM analysis showing similar structural features as *Ct*Rca1 (f). Scale bar, 50 nm.

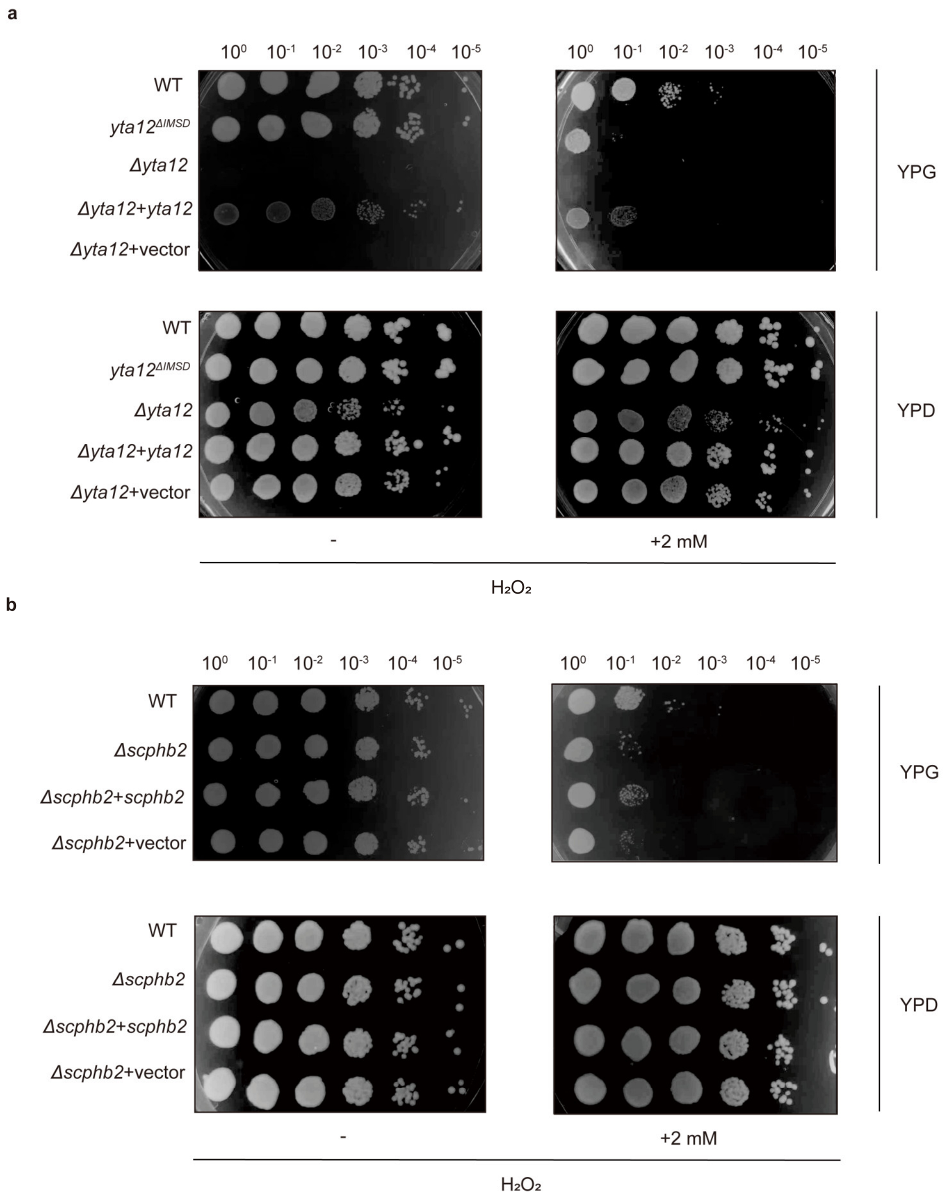


#### Figure S10. Extended data of oxidative stress sensitivity assay

**a, b,** Serial dilutions ($10^0$-$10^{-5}$) of indicated strains were spotted on YPG (respiratory) or YPD (fermentative) media ± 2 mM $H_2O_2$. Wild-type, *yta12*$^{\Delta IMSD}$ and *Δyta12* mutants with indicated complementation (a). Wild-type and *Δscphb2* strains with indicated complementation (b). Images were taken after 3 d (YPG) or 2 d (YPD) at 30°C.

**Table S1 Data collection and model validation statistics**

| | *Ct*PHB complex |
|---|---|
| **Data collection and processing** | |
| Nominal magnification | 130,000× |
| Voltage (kV) | 300 |
| Electron exposure ($e^{-}$/Å$^{2}$) | 52.4 |
| Defocus range (μm) | -1.2 to -1.8 |
| Pixel size (Å) | 1.057 |
| Micrographs | 2,900 |
| Symmetry imposed | C11 |
| Final particle images (total) | 10,623 |
| Map resolution (total) (Å) | 3.17 |
| FSC threshold | 0.143 |
| **Refinement** | |
| Model composition | |
| Chains | 22 |
| Non-hydrogen atoms | 44,715 |
| Protein residues | 5,709 |
| Nucleotides | 0 |
| Ligands | 0 |
| R.m.s. deviations | |
| Bond lengths (Å) | 0.003 |
| Bond angles (°) | 0.543 |
| Validation | |
| MolProbity score | 2.08 |
| Clashscore | 8.68 |
| Poor rotamers (%) | 4.21 |
| Ramachandran plot | |
| Favored (%) | 97.53 |
| Allowed (%) | 2.47 |
| Disallowed (%) | 0.00 |
